# De-novo genome assembly of the invasive mosquito species *Aedes japonicus* and *Aedes koreicus*

**DOI:** 10.1101/2023.04.12.534102

**Authors:** Paolo L. Catapano, Monica Falcinelli, Claudia Damiani, Alessia Cappelli, Despoina Koukouli, Paolo Rossi, Irene Ricci, Valerio Napolioni, Guido Favia

## Abstract

Recently, two invasive *Aedes* mosquito species, *Ae. japonicus* and *Ae. koreicus*, are circulating in several European countries posing potential health risks to humans and animals. Vector control is the main option to prevent mosquito-borne diseases, and an accurate genome sequence of these mosquitoes is essential to better understand their biology and to develop effective control strategies. Here, we present a de novo genome assembly of the *Ae. japonicus* (Ajap1) and *Ae. koreicus* (Akor1) based on a hybrid approach that combines Oxford Nanopore long reads and Illumina short reads data. Their quality was ascertained using various metrics. Masking of repetitive elements, gene prediction and functional annotation was performed. Sequence analysis revealed a very high presence of repetitive DNA and, among others, thermal adaptation genes and insecticide-resistance genes. The RNA sequencing analysis of larvae and adults of *Ae. koreicus* and *Ae. japonicus* exposed to different temperatures revealed genes showing a thermal-dependent activation. The assembly of Akor1 and Ajap1 genomes constitutes the first updated collective knowledge of the genomes of both mosquito species, providing the possibility to understand key mechanisms of their biology such as the ability to adapt to harsh climates and to develop insecticide-resistance mechanisms.

## Introduction

In the early nineties, Europe experienced the colonisation of vast continental areas by the *Aedes albopictus* mosquito (https://www.ecdc.europa.eu/en/disease-vectors/facts/mosquito-factsheets/aedes-albopictus). This mosquito of non-European origin is now permanently resident in the continent and it has been the protagonist of some viral epidemics as vector of numerous arboviruses and heartworms, such as Chikungunya, Zika, Dengue and filariosis (https://www.ecdc.europa.eu/en/disease-vectors/facts/mosquito-factsheets/aedes-albopictus). Also, the sudden spread in Africa of *Anopheles stephensi*, a major Asian malaria vector, is generating outbreaks even in those areas where malaria was almost eradicated (Vogel, 2022; Faulde et al, 2014). Being adapted to urban life, *An. stephensi* has the potential to further spread to many urban areas across the African continent, implying an increased number of people at risk of malaria (El-Said, 2020). Conceivably, climate change may influence the adaptation of other invasive species to new environmental niches, thus colonising new areas and rapidly spreading to others. In this frame, the appearance in Europe of two *Aedes* invasive species, namely *Ae. japonicus* and *Ae. koreicus*, is not surprising. The first European advent of *Ae. japonicus* is dated 2000, in France (Schaffner et al., 2003), while *Ae. koreicus* was reported in Belgium during the 2008 (Versteirt et al., 2012). Since then, established populations of *Ae. japonicus* were detected in Belgium, Germany, Switzerland, Austria, Slovenia, Croatia, the Netherlands, Italy, Hungary, Luxembourg, and Northern Spain. *Ae. koreicus* is well established in Italy, Germany, Russia and Hungary and it was also found in Slovenia and Switzerland (Cebrian-Camison et al., 2020). Moreover, *Ae. koreicus* has been passively spreading along the European route E35 from Italy to Germany while *Ae. japonicus* has been expanding through active dispersal (Muller et al., 2020). Thus, both species show a remarkable ability to adapt to different eco-environmental and climate conditions. Nevertheless, despite their potential role in the transmission of endemic and imported pathogens, not much is known about their real vectorial capacity and their ability to adapt to specific environmental conditions and eco-ethological contexts.

To understand the basic biology of these species, more research is needed, given the possible development of control strategies. These assumptions prompted us to conduct a study aimed at sequencing the genome of these two invasive species, with a special focus to gene clusters of peculiar interest such as those responsible for insecticide resistance and thermal adaptation.

## Results

### Genome length and GC content

Using a hybrid approach that combines Oxford Nanopore long reads and Illumina short reads data, we assembled a scaffold-level version of *Ae. koreicus* and *Ae. japonicus* genomes which length has been assessed 1.24 and 1.39 Gigabase (Gb) pairs, respectively. These dimensions resemble those of other aedine such as *Ae. aegypti* and *Ae. albopictus*, which are estimated to be respectively 1.22 (Matthews et al., 2018) and 1.19-1.28 Gb pairs (Palatini et al., 2020). The GC content of the two genomes is very similar: 39.68% in *Ae. koreicus* while 39.51% in *Ae. japonicus*. Again, these metrics are comparable to those of *Ae. aegypti* (38.3%) and *Ae. albopictus* (40.4%). Genome completeness of the two species, measured using BUSCO (Benchmarking Universal Single-Copy Orthologs) (Manni et al., 2021), showed for *Ae. koreicus* a gene completeness of 91.8%, and 8% of duplicates, while for *Ae. japonicus*, it showed 92.5% gene completeness and 13.6% duplicates. Genome annotation yielded 18,647 and 18,687 genes for *Ae. koreicus* and *Ae. japonicus*, respectively. The N50 values are 190,716 for *Ae. koreicus* and 118,241 for *Ae. japonicus* (**Figure 1, Table1**). Repeatmasker detected 71% and 71.92% of the *Ae. japonicus and Ae. koreicus* genome, respectively, as repetitive DNA.

**Figure 1.**
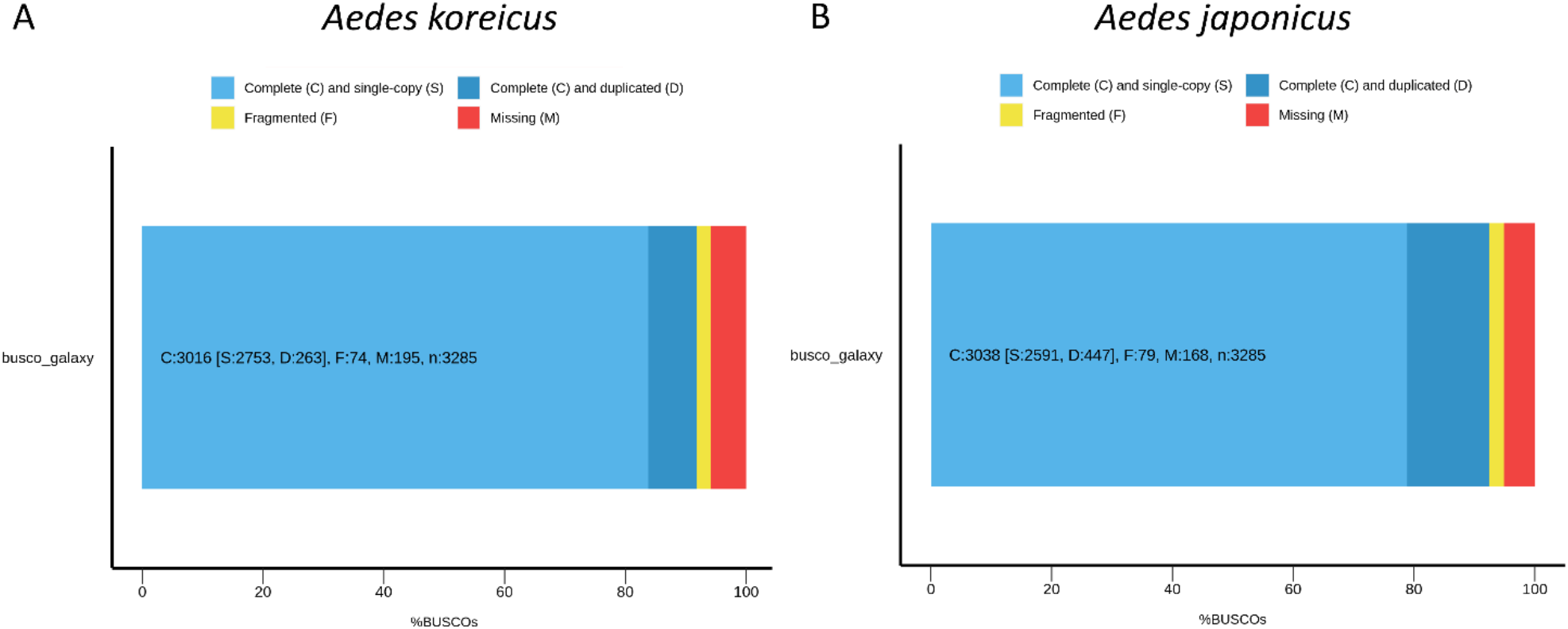
Quality assessment of the two de novo genome assembly using Benchmarking Universal Single-Copy Orthologs (BUSCO). Evaluation summary for the genome assembly of *Ae. koreicus* (A) and *Ae. Japonicus* (B) is shown with the levels of complete, fragmented, and missing gene features.

**Table 1.**
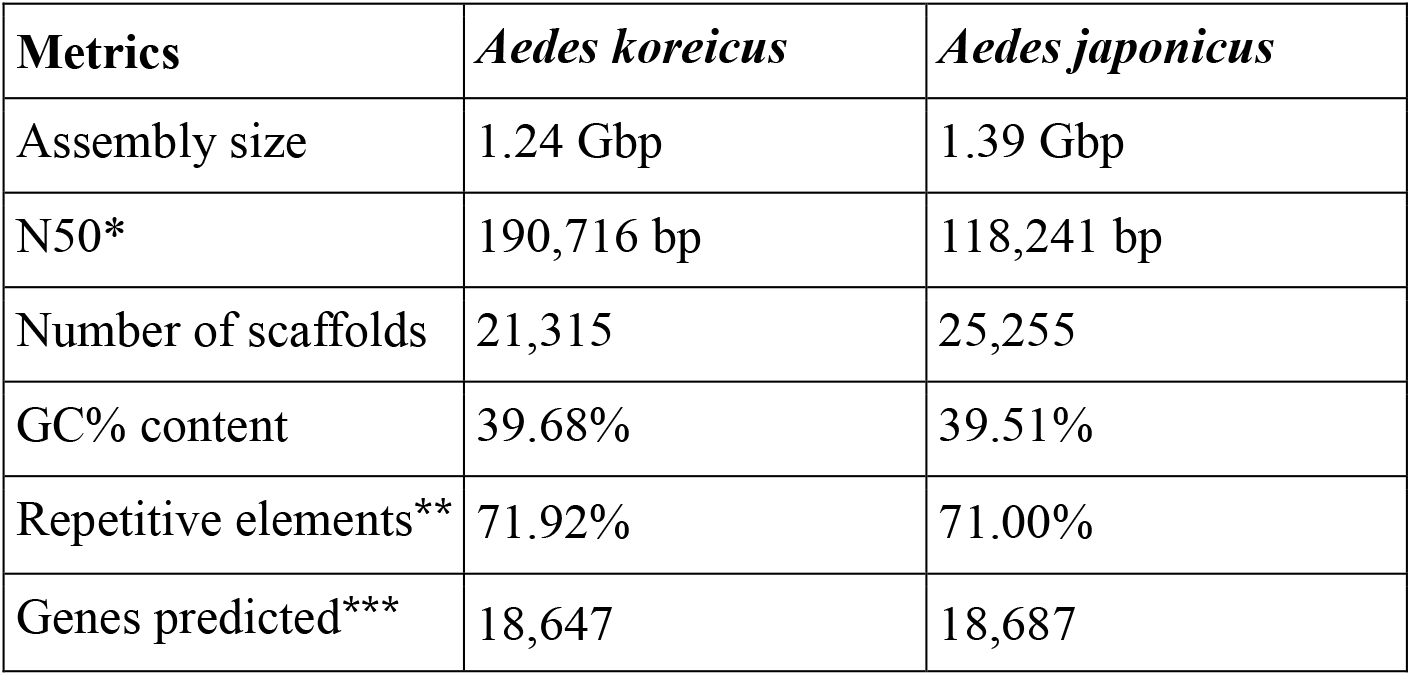
Main genome features of *Ae. koreicus* and *Ae. japonicus*. *Determined by QUAST; **Determined by RepeatMasker; *** Determined by MAKER.

| Metrics | *Aedes koreicus* | *Aedes japonicus* |
| --- | --- | --- |
| Assembly size | 1.24 Gbp | 1.39 Gbp |
| N50* | 190,716 bp | 118,241 bp |
| Number of scaffolds | 21,315 | 25,255 |
| GC% content | 39.68% | 39.51% |
| Repetitive elements** | 71.92% | 71.00% |
| Genes predicted*** | 18,647 | 18,687 |

### Phylogenetic analysis

The phylogenetic analysis was based on the comparison between the predicted proteomes of *Ae. koreicus* and *Ae. japonicus* and the proteomes of seven mosquito species belonging to three different genera (*Anopheles gambiae, Anopheles coluzzii, Anopheles arabiensis, Anopheles darlingi, Aedes aegypti, Aedes albopictus and Culex quinquefasciatus*). As expected, phylogenetic clustering revealed two main Clades. Clade I contains the four *Anopheles* species while Clade II contains the four *Aedes* species and *Cx. quinquefasciatus*. Notably, the phylogenetic tree showed that the new invasive species, *Ae. koreicus* and *Ae. japonicus*, share a most common recent ancestor (MCRA). It is worth noting that the MCRA of *Ae. koreicus* and *Ae. japonicus* diverged more recently compared to the MCRA of *Ae. aegypti* and *Ae. albopictus* (**Figure 2**).

**Figure 2.**
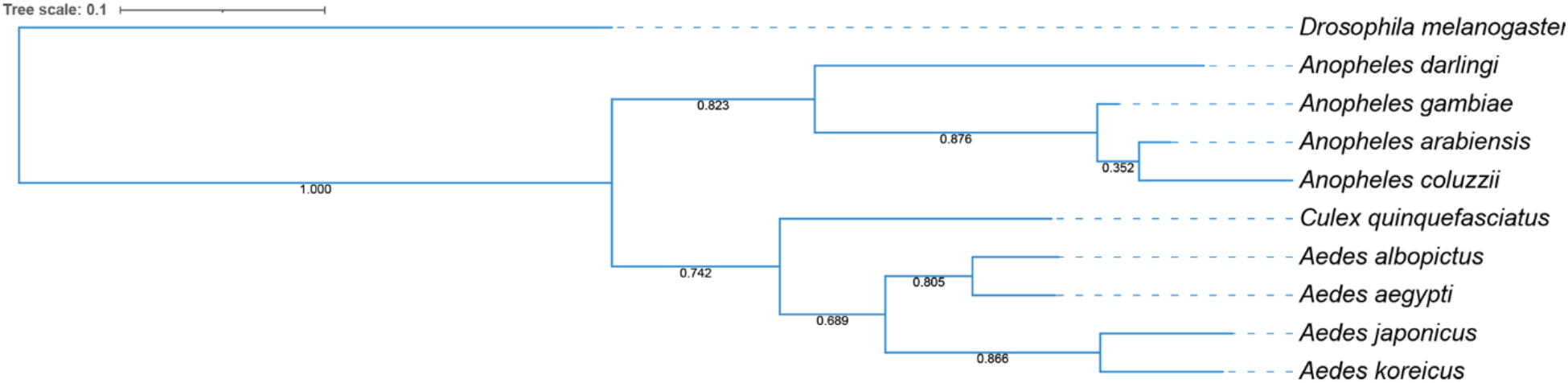
Phylogenetic analysis of the newly assembled *Ae. koreicus* and *Ae. japonicus* genomes in comparison to known mosquito species. *Drosophila melanogaster* was used as outgroup. Bootstrap values are shown at each branch.

### Annotation of specific genes

Among the several different classes of genes, particular attention has been devoted to two specific classes of genes: the ones involved in thermal adaptation and those involved in insecticide resistance.

### Thermal adaptation genes

The comparative analysis of genes involved in thermal stress between the two invasive species was extended also to *Ae. albopictus* and *Ae. aegypti*, using *Drosophila melanogaster* as reference organism (Herrmann et al., 2021). Indeed, out of 694 selected genes in *Drosophila*, we have identified 348 homologous genes in *Ae. koreicus* and 438 genes in *Ae. japonicus*. Of these, 13 are specific to *Ae. koreicus* and 35 to *Ae. japonicus*. Additional 22 genes are shared between these two species but not with *Ae. aegypti* and *Ae. albopictus* (**Supplementary Data 1**) (**Figure 3A**). Particularly, the 13 specific genes of *Ae. koreicus* are enriched in KEGG pathways involved in Pentose phosphate pathway, Fructose, mannose, galactose metabolism, Neuroactive ligand-receptor interaction, RNA degradation and Glycolysis/Gluconeogenesis. The 35 specific genes of *Ae. japonicus* are enriched in KEGG pathways such as Glycerophospholipid metabolism and Aminoacyl-tRNA biosynthesis. The 22 genes shared between these two species were enriched for KEGG pathways involved in Tryptophan, Phenylanaline, Tyrosine and Pyruvate metabolism (**Supplementary Data 2**).

**Figure 3.**
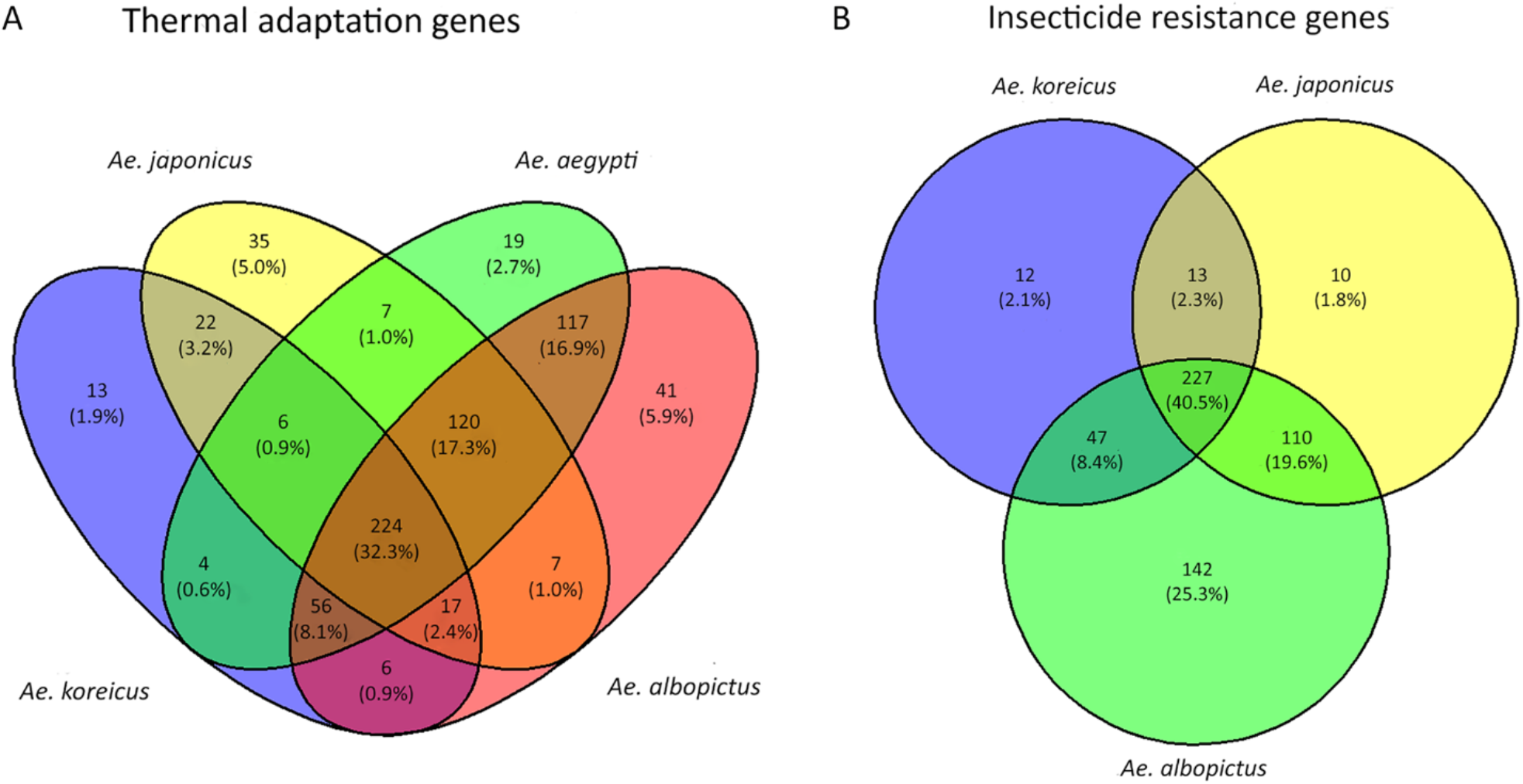
Convergence analysis of genes involved in thermal adaptation (A) and insecticide resistance (B). Venn diagrams show species-specific (unique) or shared genes among *Ae. koreicus, Ae. japonicus, Ae. aegypti* and *Ae. albopictus* mosquitoes. Thermal adaptation genes refer to *Drosophila melanogaster* and detected by homology-based search in the mosquito genomes. Insecticide resistance genes refer to *Ae. aegypti* and detected with BLAST in the genomes of *Ae. koreicus, Ae. japonicus* and *Ae. albopictus*.

### Insecticide resistance genes

The comparative analysis of genes involved in insecticide resistance between the two invasive species extended also to *Ae. albopictus* using *Ae. aegypti* as reference organism (Faucon et al., 2015). Out of 561 genes, we have identified 299 genes in *Ae. koreicus* and 360 genes in *Ae. japonicus*. Of these 12 are specific to *Ae. koreicus* and 10 to *Ae. japonicus*. An additional 13 are shared between these two species but not with *Ae. albopictus* (**Supplementary Data 3**) (**Figure 3B**). All these genes belong to gene families known to be involved in lower insecticide penetration, sequestration, and biodegradation: genes encoding for detoxification enzymes (P450, CCEs, GSTs, UGTs), cuticle proteins, ATP-binding cassette (ABC) transporters, neurotransmitter receptors, and voltage-gated channels. All three species share most of these families, nevertheless the differential distribution of members of the individual families characterizes the relative composition and quantities. More in detail: 4 out of 12 of the specific genes of *Ae. koreicus* belong to “cuticle gene family” (Chitin synthase, Pupal cuticle protein putative, Pupal cuticle protein E78 putative, Cuticle protein putative), 3 to “ion channel gene family” (Sodium leak channel non-selective protein, Voltage-dependent L-type calcium channel subunit alpha, Voltage-gated potassium channel), 2 to “ABC transporter gene family” (ATP-binding cassette sub-family A member 3, multidrug resistance protein 2/ATP-binding cassette protein c), while 2 were classified as “other detox genes” (Short-chain dehydrogenase) and one more as “other gene” (Modifier of *mdg4*).

In *Ae. japonicus* we specifically identified: one member of the “ABC transporter family” (Multidrug resistance protein 2/ATP-binding cassette protein c); one member of the “cuticle family” (Pupal cuticle protein, putative), two are ion channels (Voltage-gated potassium channel, Glutamate-gated chloride channel), two P450 gene (Cytochrome P450) and the last four remaining genes were categorized three as “other detox” (Sterol desaturase, NAD(P)H oxidase (H(2)O(2)-forming, Oxidoreductase) and one as “other” (BTB domain-containing protein). Thirteen genes are specifically shared only by *Ae. koreicus* and *Ae. japonicus*; one is an ABC transporter (ABC transporter), three belong to cuticle family (Brain chitinase and chia, Chitin synthase, Pupal cuticle protein putative), two are ion channels (Voltage-dependent L-type calcium channel subunit alpha), one is a P450 gene (Cytochrome P450). Finally, 3 out of 13 shared genes are synaptic receptors (Nicotinic acetylcholine receptor, putative, Nicotinic acetylcholine receptor beta-2 subunit putative), two are oxidoreductase (Heme peroxidase, Thioredoxin reductase) and one belong to the “other detox” family (Sterol desaturase).

### RNAseq analysis of thermal adaptation genes

Through the RNAseq analysis of larvae and adults of *Ae. koreicus* and *Ae. japonicus* exposed to different temperatures (15 and 28°C) we identified genes showing a differential temperature-dependent activation (**Figure 4**). In detail, under the low-temperature stress of 15°C several genes were differentially expressed, and the relative encoded proteins identified: in *Ae. koreicus* larvae, out of 57 upregulated and 40 downregulated transcripts, we identified 8 upregulated and 4 downregulated proteins. In adults, among 225 upregulated and 26 downregulated transcripts, we identified 27 upregulated and 2 downregulated proteins (**Supplementary Data 4**). Larvae and adults share only one upregulated gene (*aef1*, adult enhancer factor 1) but none of the downregulated (**Table 2**). In *Ae. japonicus* larvae, out of 325 upregulated and 101 downregulated transcripts, we identified 373 upregulated and 14 downregulated proteins. In adults, among 502 upregulated and 79 downregulated transcripts, we identified 70 upregulated and 24 downregulated proteins. Larvae and adults share 18 upregulated genes and 3 of the downregulated (**Table 2**). Functional GO terms were enriched in 8 Molecular functions, 1 Cellular component in larvae of *Ae. koreicus* while 2 Molecular functions in adults. Instead in *Ae. japonicus* larvae 3 Biological Processes and 1 Molecular function are enriched, while no enrichment was present in adults of *Ae. japonicus* (**Supplementary Data 5**). Notably, several differentially expressed genes seem strongly involved in thermal adaptation as they plausibly encode for specific proteins: in larvae of *Ae. koreicus*, among others of interest, we found upregulated genes encoding for Alanine/Arginine aminopeptidase (up-regulated also in adults), Acyl-CoA desaturase and Serine carboxypeptidase. Aminopeptidase activities are detected in the midgut of mosquitoes both in larvae and adults (Jakhar and Gakhar, 2019). The role of alanine aminopeptidase and more in general of aminopeptidase in thermal adaptation has been reported in different biological systems. In striped hamsters acclimated to cold (5°C), alanine aminopeptidase activity was higher than in those exposed to hot temperatures (31°C) (Zhao et al., 2014), while it is well known that psychrophilic marine bacteria produce a cold-adapted aminopeptidase (Huston et al., 2008). Concerning Acyl-CoA desaturase, there is large evidence of its involvement in cold adaptation processes: in the winged midge *Parochlus steinenii*, the extended acyl-CoA delta desaturase gene family underwent gene family expansion via multiple gene duplications for adaptation to the cold environments (Kim et al., 2022); in *Drosophila* and in Lepidoptera, the specific expression of some desaturases modulates cold adaptation mechanisms (Suito et al., 2020; Min et al., 2017). Serine carboxypeptidases are expressed in mud crabs in response to cold exposition (Chen et al., 2022).

**Table 2.** Genes differentially expressed in larvae and adults of *Ae. japonicus* and *Ae. koreicus* reared at 15°C.

| Shared genes in larvae and adults of *Aedes japonicus* |  |  |  |  |  |
| --- | --- | --- | --- | --- | --- |
| Gene Symbol | Protein name | Log2FC larvae | p-adjusted larvae | Log2FC adults | p-adjusted adults |
| *aapo1* | Alanine/arginine aminopeptidase | 11.656 | 2.790e-07 | 10.181 | 4.440e-02 |
| *abaq* | Quinolone resistance transporter | 12.385 | 2.790e-08 | 13.691 | 8.400e-03 |
| *aca13* | Putative calcium-transporting ATPase 13, plasma membrane-type | 9.773 | 1.510e-04 | 11.425 | 4.970e-02 |
| *acbp4* | Acyl-CoA-binding domain-containing protein 4 | 10.950 | 2.947e-06 | 9.561 | 4.890e-02 |
| *acoB* | Acetoin:2,6-dichlorophenolindophenol oxidoreductase subunit beta | 12.634 | 1.826e-09 | 12.130 | 4.190e-02 |
| *acr* | Acrosin | 12.487 | 2.510e-08 | 8.841 | 4.930e-02 |
| *adhT* | Alcohol dehydrogenase | 11.809 | 3.403e-08 | 11.759 | 4.620e-02 |
| *aef1* | Adult enhancer factor 1 | 5.163 | 1.248e-02 | 10.375 | 3.780e-02 |
| *aes* | Amino-terminal enhancer of split | 12.574 | 2.158e-08 | 10.135 | 4.100e-02 |
| *afg2* | ATPase family gene 2 protein | 11.695 | 2.670e-06 | 11.511 | 4.870e-02 |
| *aglA* | Probable alpha-glucosidase | 12.723 | 4.090e-09 | 12.255 | 2.950e-02 |
| *aldh9a1a* | Aldehyde dehydrogenase family | 8.228 | 7.420e-03 | 12.342 | 4.090e-02 |
| *aspC* | Aspartate aminotransferase | 12.723 | 4.090e-09 | 10.232 | 3.480e-02 |
| *baiA* | 3alpha-hydroxysteroid dehydrogenase | 10.742 | 1.159e-05 | 10.163 | 4.090e-02 |
| *cyp72a613* | Cytochrome P450 CYP72A613 | 6.693 | 4.920e-03 | 11.863 | 4.510e-02 |
| *cyp72a616* | Cytochrome P450 CYP72A616 | 12.842 | 8.839e-09 | 11.863 | 4.510e-02 |
| *ksdD* | 3-oxosteroid 1-dehydrogenase | 9.216 | 1.220e-03 | 13.266 | 3.190e-02 |
| *acr* | Acrosin | -5.629 | 1.300e-03 | -10.732 | 3.229e-02 |
| *aef1* | Adult enhancer factor 1 | -8.494 | 3.000e-04 | -9.639 | 4.840e-02 |
| *ab* | Protein abrupt | -12.959 | 1.129e-10 | -9.678 | 4.710e-02 |
| Shared genes in larvae and adults of *Aedes koreicus* |  |  |  |  |  |
| Gene Symbol | Protein name | Log2FC larvae | p-adjusted larvae | Log2FC adults | p-adjusted adults |
| *aef1* | Adult enhancer factor 1 | 20.775 | 1.580e-04 | 20.580 | 3.287e-06 |

**Figure 4.**
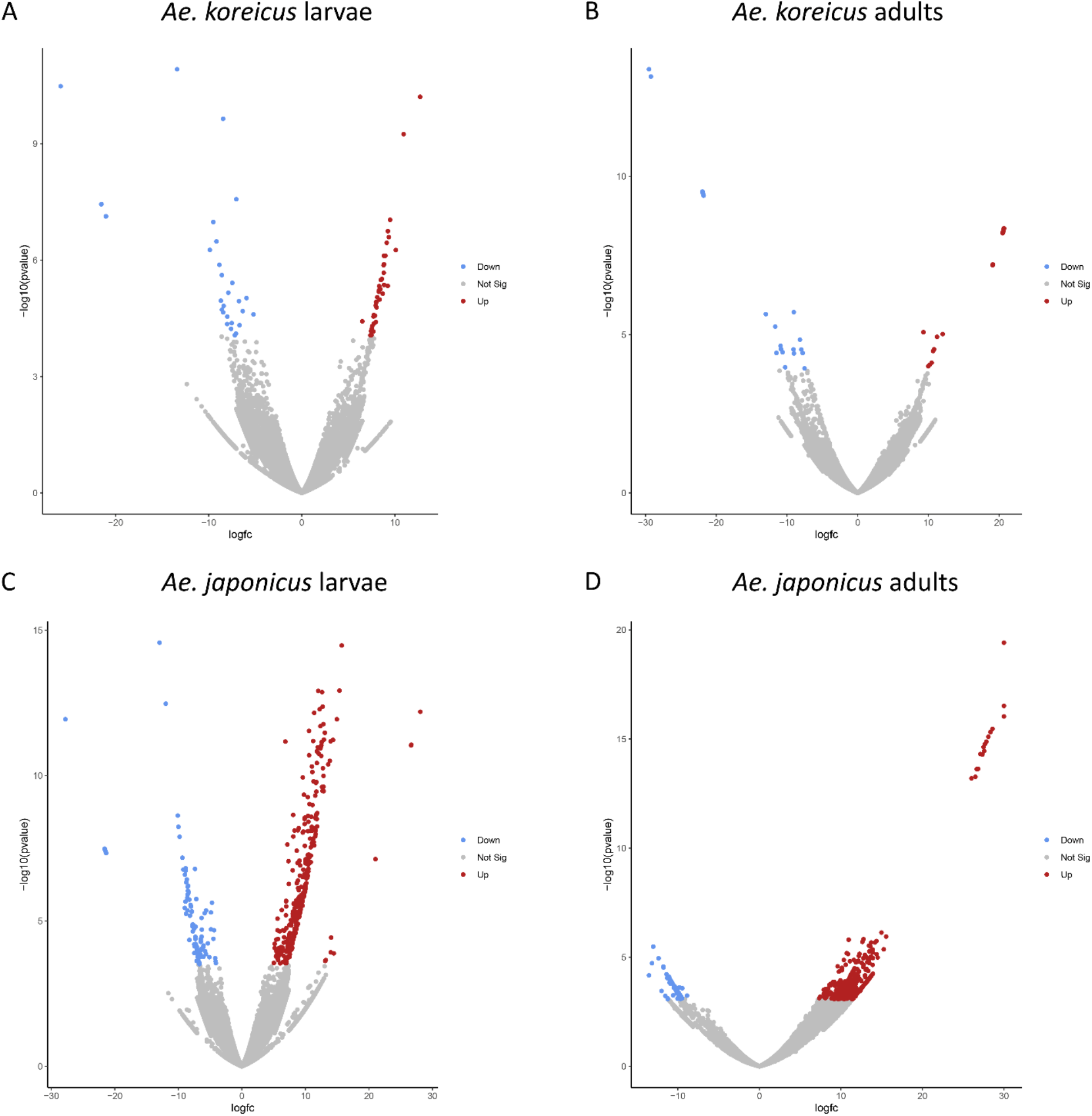
Analysis of differentially expressed genes in larvae and adults of *Ae. koreicus* and *Ae. japonicus* following thermal adaptation. Volcano plots comparing transcripts obtained by samples reared at 15°C and 28°C (control group): upregulated (red), downregulated (blue) and not significantly altered (grey) transcripts are shown in *Ae. koreicus* larvae (A), *Ae. koreicus* adults (B), *Ae. japonicus* larvae (C) and *Ae. japonicus* adults (D) respectively.

In adults of *Ae. koreicus*, among many other genes likely involved in thermal regulation, we found up-regulated genes encoding for Cytochrome P450, Fatty acyl-CoA reductase, and Mitochondrial adenine nucleotide transporter/translocase. Insect’s cytochrome P450 is well known to be involved in tolerance mechanisms (Franke et al., 2019; Huang et al., 2017) and in some insects, as the Crysomelidae *Galeurca daurica*, is upregulated together with a Fatty acyl-CoA reductase during recovery from cold stress (Zhou et al., 2019). Mitochondrial adenine nucleotide transporter/translocase has roles in thermal adaptation in several organisms as for example in thermal acclimation in the rainbow trout *Oncorhyncus mykiss* (Kraffe et al., 2007), or in response to thermal stress in *Apostichopus japonicus* (Liu et al., 2016).

Among the genes downregulated in larvae of *Ae. koreicus*, after exposure to 4 °C, we found a gene encoding a 5’-AMP-activated protein kinase (catalytic subunit alpha-2); interestingly activation of an AMP-activated protein kinase in response to temperature elevation has been reported in the zebra mussel, *Dreissena polymorpha* (Jost et al., 2015)

Two genes were found downregulated in adults of *Ae. koreicus* when exposed to cold conditions: an Acetyl-CoA carboxylase (*acc*) and a Steroid receptor seven-up, isoform A. *acc* is reported to have a role in thermal adaptation in sheep, more precisely its activity seems depressed in tissues exposed to cold (Moibi et al., 2000). The Steroid receptor seven-up is involved in *Drosophila* oogenesis, thus influencing final egg output (Weaver and Drummond Barbosa, 2019).

In *Ae. japonicus* larvae, among the many up-regulated genes, plausibly involved in the mechanisms of cold adaptation after exposure to 15°C, there are some common to those previously described *in Ae. koreicus* (e.g., Cytochrome P450). Among the many others of interest are those encoding some dehydrogenases (both alcohol and 6-phosphogluconate dehydrogenase), a multicopper oxidase, a glycosyl hydrolase and helicases. Indeed, in *Bactrocera dorsalis* low-temperature stress induced higher alcohol dehydrogenase activities in different life developmental stages with higher increase intensity in adults and pupae than in larvae (Wang et al., 2014). 6-phosphogluconate dehydrogenase it is known to play a role in increasing cold tolerance in some plants (Tian et al., 2021; Sardesai et al., 2001).

Multicopper oxidases as well are involved in mechanisms of cold tolerance in different organisms as plant and microbes (Xu et al., 2022; Karmacharya et al., 2022), while glycosyl hydrolases contribute to cold adaptation in many different organisms ranging from many plants to some yeasts up to the Antarctic springtail (Song et al., 2017). Some members of the helicase family behave similarly with different plants and algae (Peng et al., 2021; Wang et al., 2020).

In adults of *Ae. japonicus*, among the many genes up-regulated after exposure to 15°C, we should mention the members of the Cytochrome P450 and Acyl-CoA desaturase families which role in cold adaptation has already been described (Iqbal et al., 2022). Moreover, genes of the Actin family, that in *Culex pipiens* are known to be expressed from early diapause to late diapause and in young non-diapaused adult mosquitoes reared at 18°C (Kim et al, 2006), were found up-regulated, as well as members of Mitochondrial carriers and members of the Methyltransferase superfamily. Interestingly in some tick’s species, DNA methyltransferase are known to contribute to cold tolerance (Agwunobi et al., 2021). Other genes found up-regulated are members of the aminotransferase family that, as shown in Corn Borer *Ostrinia nubilalis*, during diapause and cold hardening catalyse the production of L-alanine, an important cryoprotectant (Uzelac et al, 2020).

In both *Ae. japonicus* larvae and adults the gene encoding the adult enhancing factor 1 (*aef1*) was found down regulated after exposure to 15 °C. Among other functions, this factor has been shown to bind the alcohol dehydrogenase adult enhancer site (AAE) thus regulating its transcription (Potter et al., 1994). In adults, as already seen in *Ae. koreicus*, a gene encoding a 5’-AMP-activated protein kinase catalytic sub-unit alpha was also downregulated when exposed to cold.

## Discussion

Mosquito invasive species, having the potential to transmit a range of different pathogens, pose significant public health problems where they establish, and their ranges and potential impacts are shifting with climate change. This is well demonstrated by the thirty-year experience on the stabilization of *Ae. albopictus* in Europe and, more recently, on the impact of the transmission of urban malaria in relation to the arrival of *An. stephensi* in Africa (Vogel, 2022). Although some information about genome organisation has been very partially provided for *Ae. koreicus* (Kurucz et al., 2022), a more detailed knowledge of the basic biology of the new European invasive species *Ae. koreicus* and *Ae. japonicus* is a fundamental prerequisite to control these insect vectors. Hence, genome sequencing of both species may provide insight into the genetics basis of their competence for pathogens transmission and for the development of species-specific control methods.

Both species are characterized by a genome size and GC content comparable to other aedines genomes (*Ae. albopictus* and *Ae. aegypti*). Nevertheless, the two species are phylogenetically strictly correlated in respect to all the other mosquito species taken in consideration. This suggests common mechanisms of adaptation to eco-ethological contexts and could explain their almost contemporary appearance in large areas of Mediterranean and central Europe.

Consequently, forecasting changes expanding the regions that are suitable to invasion of *Aedes* vectors and *Aedes*-borne viruses, with regards to dengue, chikungunya, and Zika, is a key element of public health preparedness. Moreover, the insecticide resistance developed by several mosquito vectors are undermining the effectiveness of their control. Thus, we focused on those genes implicated in both thermal adaptation and insecticide resistance. The analysis of both group of genes revealed some intriguing features. Both *Ae. koreicus* and *Ae. japonicus* are characterized by species-specific set of genes as well as genes that are shared between these two species but not by other aedines. Considering the function(s) of the proteins encoded by these genes (e.g., decarboxylases) it is very likely that most, if not all, of these genes drive the specific behaviour of the two mosquito species in climatic adaptation and insecticide resistance. This is further substantiated by the RNA-seq analysis following exposure of larvae and adults to different temperatures. Indeed, as reported in the results, the expression of several genes has shown to be strongly modulated by the temperature and some of these genes would seem to be involved in the adaptation to low temperatures and could, consequently, contribute on the one hand to a better understanding of the mechanisms underlying the geographical distributions of the two invasive species, on the other to better monitor and control the dispersion of the two species. The ability to monitor and control vector mosquitoes is also supported by the ability to use insecticides and biocides wisely in relation to the possible onset of insecticide-resistance. In this frame, the identification of genes plausibly involved in possible insecticide resistance mechanisms (e.g., Chitin synthases), can represent an excellent basis on which to build further monitoring and control programmes.

Ultimately, despite the need for future corroborating studies, the sequencing of the two genomes and the analysis of the two selected groups of genes pave the way to the possibility of specific control strategies aimed to limit the risks associated with the recent introduction of the two invasive species in Europe.

## METHODS

### Collection of Samples

Larvae of *Ae. koreicus* and *Ae. japonicus* were collected in two villages situated in Veneto region (North-east Italy), Alano di Piave (45°54’26’’ N, 11°54’28’’ E), and Feltre (46°0’49.903”N 11°53’49.996”E) respectively. Larvae were mainly collected from artificial containers by dipping and delivered to the insectary of Camerino University. Four instar larvae were morphologically identified according to Montarsi et al., 2013. The DNAs of newly emerged adults were used for the genome analysis. The RNA of four instar larvae and adults were used for the RNA sequencing analysis.

Total genomic DNA were obtained from pools of three females of *Ae. koreicus* and *Ae. japonicus*. DNA was extracted using a JetFlex Genomic DNA Purification kit (Invitrogen, Thermo Fisher Scientific, Waltham, MA, USA) according to the manufacturer’s instructions.

For the RNA seq analysis samples were prepared from two different cohorts of adults and four instar larvae *Ae. koreicus* and *Ae. japonicus* reared at 15°C and 28°C respectively. Total RNAs were extracted from single adult and pool of five four instar larvae using RNAzol reagent (Sigma-Aldrich USA), according to the manufacturer’s instructions.

### Sequencing

Samples of *Ae. koreicus* and *Ae. japonicus* were sent to IGAtech (Udine, Italy) for short and long read sequencing. Briefly, for short read sequencing: CeleroTM DNA-Seq kit (NuGEN, San Carlos, CA) has been used for library preparation following the manufacturer’s instructions and was sequenced with NovaSeq 6000 in paired-end 150 bp mode. Long read sequencing was performed with Oxford Nanopore PromethION. The samples were prepared using the kit SQK-LSK109 and sequenced on a flowcell R9.4.1. Basecalling of data was performed with Guppy 5.0.13. Reads were filtered with minimum qscore of 7 and length of 500.

For RNA-seq sequencing, Universal Plus mRNA-Seq kit (Tecan Genomics, Redwood City, CA) has been used for library preparation. Libraries were then prepared for sequencing and sequenced on paired-end 150 bp mode on NovaSeq 6000 (Illumina, San Diego, CA).

The total reads count generated by NovaSeq 6000, and Oxford Nanopore PromethION are reported in the **Supplementary Data 6**.

### Assembly

The Illumina reads were error-corrected using BFC release 181 (Li, 2015). The Nanopore reads were error-corrected using LoRDEC v0.9 (Salmela and Rivals, 2014) with the error-corrected Illumina overlapping Paired-End (PE) reads, a k-mer size of 19 and a solidity threshold of 3. A first assembly was performed using FLYE 2.9.1 (Kolmogorov et al., 2019) with the long raw Oxford Nanopore reads. The resulting assembly was polished using two rounds of HyPo v1.0.3 (Kundu et al., 2019) with the error-corrected Illumina overlapping PE reads. Scaffolding was then performed with two rounds of LINKS v2.0.1(Warren et al., 2015) and ntLINKS v1.3.4 (Coombe et al., 2021) using the gap-filling option and with the error-corrected Illumina PE and Nanopore libraries. An additional round of HyPo, as previously described, was performed. We removed haplotig contamination by using “purged dups (Guan et al., 2020) to produce the final assembly. The assembly quality was assessed by computing two metrics: QUAST v5.0.2 (Mikheenko et al, 2018) and BUSCO v4.0.5 (Manni et al, 2021), using as the lineage Diptera.

### RNA sequencing

Adapter sequences and low-quality bases were trimmed (Trimmomatic) (Bolger, 2014) and quality checks before and after trimming were performed (Fastqc) (Andrews, 2010). De novo Assemblies were done with SPAdes (Bushmanova et al., 2019), for RNA. Quality of assemblies was assessed with BUSCO (Manni et al., 2021) and QUAST (Mikheenko et al., 2018). Error-corrected reads were mapped to transcripts and read count was performed with SALMON (Patro et al., 2017). Read count files were then used to compare gene expression of two groups, cold (15°C) and hot (28°C) using DESeq2 pipeline in RStudio (Love et al., 2014). We performed transcript annotation using Trinotate pipeline (Bryant et al., 2017). Gene ontology and functional annotation was done with UniProt (The UniProt Consortium, 2023) and Enrichr (Chen et al., 2013).

### Annotation

The automatic annotation of the genome was performed using the MAKER pipeline (Holt and Yandell, 2011). We provided MAKER data with RNA-seq data of *Ae. koreicus* and *Ae. japonicus* sequenced by our lab and additonal ESTs and RNA-seq data from similar mosquito species such as *Ae. aegypti, Ae. albopictus, Cx. quinquefasciatus* that were publicly available. To predict gene models SNAP (Korf, 2004) and Augustus (Stanke and Morgenstern, 2006) within MAKER were used. To detect repetitive elements in the genomes, we used RepeatMasker (Tarailo-Graovac and Nansheng, 2009). Functional annotation was performed with the InterProScan 5 Pipeline (Jones et al., 2014).

### Phylogenetic analysis

Comparative genomic analysis was performed with the Orthofinder Pipeline (Emms and Kelly, 2015). The two new assemblies of *Ae. koreicus* and *Ae. japonicus* were compared with mosquito genomes previously assembled like *Ae. aegypti, Ae. albopictus, Cx. quinquefasciatus, An. gambiae, An. coluzzii, An. arabiensis, An. darlingi* and using *D. melanogaster* genome as an outgroup. The phylogenetic tree graph was modified using iTOL v6 (Letunic and Bork, 2021).

Detection of cold tolerance genes and insecticide resistance genes in the new assemblies For the detection of the cold tolerance genes, protein sequences of genes found to be differentially expressed in D. melanogaster at cold temperatures compared to warmer temperatures (Herrmann and Yampolsky, 2021) were mapped with TBLASTN (Gertz et al., 2006) to the assemblies of *Ae. koreicus, Ae. japonicus, Ae. albopictus* and *Ae. aegypti*. We filtered results based on hit scoring (>200), and in the case of multiple hits in the same genomic location, only the hit with the highest score was maintained. To find the genes specific to the different mosquito species, we compared the results and built Venn diagrams with RStudio. Functional annotation enrichment analysis was performed with Flyenrichr (Chen et al., 2013).

For the detection of insecticide resistance genes in the new assemblies, we mapped the protein sequences of 751 metabolic insecticide resistance genes of *Ae. aegypti* (Faucon, 2015) using TBLASTN (Gertz et al., 2006), against the assemblies of *Ae. koreicus, Ae. japonicus* and *Ae. albopictus*. We filtered results based on hit scoring (>200), and in the case two hits occurred in the same genomic location, we kept the hit with the highest score. We then compared results between species to detect genes specific to *Ae. koreicus* and *Ae. japonicus* and not present in *Ae. albopictus*. Venn diagrams were built with RStudio.

## Data availability

All the reads related to the mosquito genome sequencing and RNA sequencing of *Ae. koreicus* and *Ae. japonicus* described in this study have been deposited in the NCBI with accession numbers PRJNA947548 and PRJNA947978, respectively.

## Acknowledgements

**…**.

