## Supplementary table 1 for "De-novo genome assembly of the invasive mosquito species *Aedes japonicus* and *Aedes koreicus*"

**Supplementary data 1A**

**Thermal adaptation genes found in *Aedes koreicus***

NP_001137782.1
NP_001262724.1
NP_001286496.1
NP_001285616.1
NP_001285121.1
NP_001245529.1
NP_001286607.1
NP_788494.1
NP_524264.1
NP_001137779.1
NP_001096908.1
NP_001014584.1
NP_001188892.1
NP_001163722.1
NP_001137706.1
NP_001163731.1
NP_001260120.1
NP_001262509.1
NP_001245448.1
NP_611218.1
NP_572981.1
NP_476817.1
NP_524337.1
NP_001188751.1
NP_649732.1
NP_001137750.1
NP_724679.2
NP_001245791.1
NP_001246581.1
NP_001163649.1
NP_651960.2
NP_647937.2
NP_001097888.1
NP_524470.4
NP_651004.1
NP_001097880.1
NP_995670.2
NP_001286949.1
NP_001163024.1
NP_001097409.1
NP_001285580.1
NP_001260177.1
NP_650836.1
NP_001287316.1
NP_001247197.1
NP_523800.1
NP_611386.2
NP_001287443.1
NP_001036351.2
NP_523578.1
NP_001246073.1
NP_001260023.1
NP_001036454.1

NP_001097423.1
NP_611584.1
NP_001097378.1
NP_001303433.1
NP_001262793.1
NP_650475.1
NP_001285176.1
NP_996408.1
NP_001188784.1
NP_001189278.1
NP_609496.1
NP_001285090.1
NP_001285408.1
NP_523613.4
NP_001097982.1
NP_610446.1
NP_001287044.1
NP_001246914.1
NP_001261026.1
NP_523868.1
NP_001246140.1
NP_001261434.1
NP_001014720.1
NP_648053.1
NP_608781.2
NP_732407.1
NP_001036362.1
NP_523581.1
NP_476899.1
NP_001036691.1
NP_650223.1
NP_001097517.1
NP_610107.1
NP_728468.1
NP_649225.1
NP_477262.1
NP_650963.2
NP_001097889.2
NP_001260111.1
NP_001097922.1
NP_651515.1
NP_996221.1
NP_001260679.1
NP_608557.4
NP_650029.1
NP_732900.1
NP_001246699.1
NP_001286308.1
NP_648918.1
NP_525081.1
NP_001189276.1
NP_652625.2
NP_001163387.2
NP_001162804.1
NP_001014573.1
NP_001287406.1

NP_001262626.1
NP_001285859.1
NP_001027059.1
NP_649370.1
NP_001189282.1
NP_001287203.1
NP_001097740.1
NP_001285915.1
NP_001285680.1
NP_001246357.1
NP_609528.3
NP_001261088.1
NP_001189170.1
NP_001245832.1
NP_001246145.1
NP_001261474.1
NP_001097237.1
NP_001285908.1
NP_001137649.1
NP_001036269.2
NP_722744.1
NP_611140.3
NP_001287054.1
NP_648026.1
NP_001262398.1
NP_001262088.1
NP_652731.1
NP_610804.1
NP_001033881.1
NP_001262652.1
NP_610121.2
NP_648389.3
NP_001261618.1
NP_608968.2
NP_001260667.1
NP_649008.2
NP_001246037.1
NP_001260974.1
NP_001286637.1
NP_001188754.1
NP_001245916.1
NP_001285505.1
NP_477307.1
NP_001163749.1
NP_729935.1
NP_524157.2
NP_651481.1
NP_001247251.1
NP_511068.2
NP_649435.2
NP_611654.1
NP_001097890.1
NP_001247313.1
NP_001261613.1
NP_477311.1
NP_001286835.1

NP_001260304.1
NP_651116.1
NP_610940.1
NP_001260616.1
NP_001246805.1
NP_001286202.1
NP_001286069.1
NP_523507.2
NP_001188761.1
NP_001286383.1
NP_611595.3
NP_609500.1
NP_647979.1
NP_001027104.1
NP_001246653.1
NP_611542.2
NP_001162795.1
NP_001259941.1
NP_731872.1
NP_001036752.1
NP_001262929.1
NP_573357.1
NP_650563.2
NP_572259.1
NP_001259573.1
NP_001285570.1
NP_001189037.1
NP_001284920.1
NP_651601.1
NP_001285354.1
NP_001246401.1
NP_001097333.1
NP_649676.1
NP_611262.1
NP_001137835.2
NP_651792.1
NP_651040.3
NP_001014475.1
NP_001027277.1
NP_001163032.1
NP_001261255.1
NP_001245889.1
NP_001262615.1
NP_001036736.2
NP_001246234.1
NP_001260487.1
NP_609119.2
NP_001261253.1
NP_001188590.1
NP_001260534.1
NP_001162875.1
NP_001303445.1
NP_001097258.2
NP_001036715.1
NP_647991.1
NP_001261543.1

NP_572382.1
NP_001285254.1
NP_001262027.1
NP_610185.1
NP_001356938.1
NP_524339.1
NP_001097533.1
NP_524074.1
NP_001097127.1
NP_001036349.2
NP_001287456.1
NP_726254.1
NP_001189101.1
NP_001097183.1
NP_001260153.1
NP_650643.2
NP_001285238.1
NP_524001.2
NP_648665.1
NP_001036757.1
NP_525117.2
NP_523450.1
NP_001097077.1
NP_001097501.1
NP_001189222.1
NP_001286506.1
NP_001259495.1
NP_001014644.3
NP_001245907.1
NP_001286665.1
NP_573392.2
NP_001014535.1
NP_523888.1
NP_610454.2
NP_001245933.1
NP_001261878.1
NP_001246206.1
NP_001162998.1
NP_001260517.1
NP_651063.1
NP_523600.5
NP_650920.3
NP_001246288.1
NP_648758.1
NP_001163646.1
NP_001284923.1
NP_524382.1
NP_648662.1
NP_572452.1
NP_001162885.1
NP_001262717.1
NP_001097148.1
NP_001162909.1
NP_001188769.1
NP_729109.1
NP_001260085.1

NP_001245914.1
NP_001097105.3
NP_788278.1
NP_477052.2
NP_611811.3
NP_001097226.1
NP_001285425.1
NP_001162922.1
NP_001259858.1
NP_001262617.1
NP_001097631.1
NP_001287128.1
NP_001262655.1
NP_001261156.1
NP_001261908.1
NP_610724.1
NP_609579.1
NP_731547.1
NP_001014732.1
NP_001286071.1
NP_001287441.1
NP_733378.1
NP_476741.1
NP_001261396.1
NP_001163667.1
NP_001260541.1
NP_001097038.2
NP_001246983.1
NP_651521.1
NP_001259851.1
NP_001261039.1
NP_611185.3
NP_611952.1
NP_610049.1
NP_001162777.1
NP_001259689.1
NP_573173.2
NP_648992.1
NP_609160.2
NP_648027.3
NP_001260841.1
NP_001189236.1
NP_651411.1
NP_995812.1
NP_650287.3
NP_996277.1
NP_001261816.1
NP_001033835.2
NP_650317.3
NP_649428.1
NP_001287390.1
NP_001285984.1
NP_001285958.1
NP_651286.3
NP_001261083.1
NP_611633.2

NP_609787.2
NP_001097322.1
NP_650773.1
NP_649511.2
NP_648509.1
NP_001014631.3
NP_001097425.1
NP_001262456.1
NP_001285274.1
NP_476831.1
NP_524938.1
NP_001287331.1
NP_001137843.1
NP_001163592.1
NP_001138101.1

**Supplementary data 1B**
**Thermal adaptation genes found in *Aedes japonicus***
NP_001262724.1
NP_001260059.1
NP_001284845.1
NP_001262953.1
NP_001262200.1
NP_001261301.1
NP_648918.1
NP_523507.2
NP_647937.2
NP_001285121.1
NP_001286607.1
NP_001261026.1
NP_001096908.1
NP_001245529.1
NP_524830.2
NP_788494.1
NP_524339.1
NP_524264.1
NP_476741.1
NP_001163749.1
NP_611140.3
NP_001137782.1
NP_001027193.2
NP_001245907.1
NP_649886.1
NP_001137706.1
NP_001285655.1

NP_001188590.1
NP_611218.1
NP_001137779.1
NP_001163731.1
NP_001260120.1
NP_648665.1
NP_001138102.1
NP_477307.1
NP_001036374.1
NP_001262509.1
NP_476899.1
NP_572981.1
NP_001260291.1
NP_001188875.1
NP_572884.1
NP_001261115.1
NP_609813.1
NP_001260350.1
NP_001096975.1
NP_001188751.1
NP_524337.1
NP_001137910.2
NP_001245791.1
NP_001262639.1
NP_001246581.1
NP_001137778.2
NP_001334713.1
NP_001036736.2
NP_001163649.1
NP_001260388.1
NP_476817.1
NP_001014584.1
NP_650004.1
NP_001096941.2
NP_001285404.1
NP_001245916.1
NP_001097888.1
NP_001285589.1
NP_651004.1
NP_649289.2
NP_651481.1
NP_001285580.1
NP_001163465.1
NP_001262324.1
NP_649026.1
NP_477262.1
NP_001286949.1
NP_001163024.1
NP_001097409.1
NP_523868.1
NP_001259194.1
NP_001027277.1

NP_996221.1
NP_001260023.1
NP_610940.1
NP_001246073.1
NP_001097072.1
NP_650836.1
NP_001246914.1
NP_001285238.1
NP_611386.2
NP_001286472.1
NP_724679.2
NP_001097183.1
NP_001188757.1
NP_001262793.1
NP_001286102.1
NP_996408.1
NP_730938.1
NP_001097745.1
NP_001189278.1
NP_001285408.1
NP_001286090.1
NP_523613.4
NP_610446.1
NP_001246140.1
NP_001137707.1
NP_001097982.1
NP_001097077.1
NP_001245941.1
NP_523581.1
NP_001036362.1
NP_609566.1
NP_609204.2
NP_476905.1
NP_732407.1
NP_001097889.2
NP_650223.1
NP_001036474.2
NP_649225.1
NP_525117.2
NP_001097922.1
NP_651515.1
NP_608557.4
NP_001033835.2
NP_001287316.1
NP_001260679.1
NP_650029.1
NP_650963.2
NP_609040.1
NP_609999.2
NP_648053.1
NP_001246315.2
NP_001368942.1

NP_001285908.1
NP_609496.1
NP_722744.1
NP_001262655.1
NP_001285915.1
NP_652625.2
NP_001162804.1
NP_610232.1
NP_523800.1
NP_001284920.1
NP_728468.1
NP_001286071.1
NP_001188892.1
NP_001285859.1
NP_001287044.1
NP_001287406.1
NP_001245448.1
NP_001246206.1
NP_001245832.1
NP_649370.1
NP_001189282.1
NP_001162795.1
NP_651792.1
NP_609528.3
NP_001287203.1
NP_001260447.1
NP_650138.1
NP_001263109.1
NP_001261999.1
NP_001246357.1
NP_001097890.1
NP_001163413.1
NP_001285096.1
NP_610804.1
NP_001246145.1
NP_524074.1
NP_001189170.1
NP_001261748.1
NP_001097880.1
NP_001285354.1
NP_651960.2
NP_001261088.1
NP_648026.1
NP_723592.1
NP_001260177.1
NP_001286094.1
NP_001138163.1
NP_001262398.1
NP_524157.2
NP_652731.1
NP_648758.1
NP_995812.1

NP_001285042.1
NP_001262652.1
NP_611654.1
NP_001260667.1
NP_525081.1
NP_729935.1
NP_610121.2
NP_001286637.1
NP_001162922.1
NP_001014572.1
NP_001285505.1
NP_001285090.1
NP_001097535.1
NP_001261411.1
NP_001261396.1
NP_524071.2
NP_647623.1
NP_001033936.1
NP_001246699.1
NP_001247197.1
NP_511068.2
NP_650886.1
NP_001260930.1
NP_001287623.1
NP_001097608.1
NP_001014642.4
NP_001097555.1
NP_649435.2
NP_001014475.1
NP_001303445.1
NP_001247313.1
NP_001286496.1
NP_001262043.1
NP_001284785.1
NP_477311.1
NP_001259089.1
NP_001188784.1
NP_648389.3
NP_001189276.1
NP_651116.1
NP_476768.1
NP_001262722.1
NP_001036700.1
NP_001163781.1
NP_001286835.1
NP_001260616.1
NP_001262316.1
NP_001286383.1
NP_477004.5
NP_001033881.1
NP_001097038.2
NP_001260304.1

NP_001286202.1
NP_001188761.1
NP_611595.3
NP_001246805.1
NP_001014732.1
NP_001137649.1
NP_001262929.1
NP_523364.2
NP_001188733.1
NP_648992.1
NP_001263066.1
NP_611542.2
NP_650563.2
NP_001036752.1
NP_573357.1
NP_572259.1
NP_001097369.2
NP_001260434.1
NP_001188909.1
NP_001014621.1
NP_651601.1
NP_001162639.2
NP_001246401.1
NP_001246351.1
NP_001097533.1
NP_650867.1
NP_611262.1
NP_649676.1
NP_001137835.2
NP_001245483.1
NP_001246034.1
NP_610235.1
NP_610107.1
NP_001285642.1
NP_001261255.1
NP_609119.2
NP_001260487.1
NP_001261618.1
NP_001262615.1
NP_001096890.1
NP_001162875.1
NP_001287361.1
NP_001260534.1
NP_524470.4
NP_001027070.1
NP_001014644.3
NP_610185.1
NP_001097258.2
NP_001262670.1
NP_572382.1
NP_001033952.1
NP_647991.1

NP_610276.1
NP_001014559.1
NP_001033855.1
NP_001284923.1
NP_476831.1
NP_001263017.1
NP_651411.1
NP_001262027.1
NP_001163667.1
NP_001097127.1
NP_001189101.1
NP_477052.2
NP_726254.1
NP_001036349.2
NP_001260153.1
NP_651841.3
NP_001097129.1
NP_732900.1
NP_001285274.1
NP_477132.1
NP_610217.1
NP_650643.2
NP_001286506.1
NP_001097556.2
NP_572775.1
NP_001036757.1
NP_609579.1
NP_573392.2
NP_001259495.1
NP_001262553.1
NP_001262557.1
NP_001247239.1
NP_650086.2
NP_649895.1
NP_001260517.1
NP_001188754.1
NP_001097707.1
NP_610318.1
NP_001260869.1
NP_523888.1
NP_001285176.1
NP_610031.1
NP_001286589.1
NP_001284892.1
NP_651063.1
NP_523600.5
NP_001246288.1
NP_611185.3
NP_001097435.2
NP_001162998.1
NP_001262088.1
NP_001285682.1

NP_001262191.1
NP_001162885.1
NP_001188915.1
NP_788714.1
NP_572452.1
NP_524382.1
NP_001262717.1
NP_001097148.1
NP_524875.1
NP_651581.2
NP_611811.3
NP_729109.1
NP_477213.1
NP_001245914.1
NP_001036635.1
NP_001097425.1
NP_001285425.1
NP_001262617.1
NP_001097631.1
NP_001287128.1
NP_001027059.1
NP_647793.1
NP_001163387.2
NP_651456.1
NP_001260541.1
NP_001137878.2
NP_001162777.1
NP_001262131.1
NP_001261908.1
NP_001287441.1
NP_731547.1
NP_610724.1
NP_001262998.1
NP_001261039.1
NP_001034006.1
NP_725114.1
NP_001188999.1
NP_648236.1
NP_001247251.1
NP_001261156.1
NP_648280.1
NP_001262614.1
NP_572297.1
NP_001246983.1
NP_611952.1
NP_001036351.2
NP_001097074.1
NP_610049.1
NP_001097261.1
NP_523801.2
NP_001014573.1
NP_001188904.1

NP_609500.1
NP_001097705.1
NP_609160.2
NP_650672.1
NP_996277.1
NP_573173.2
NP_788515.1
NP_001260974.1
NP_001036454.1
NP_001285958.1
NP_650287.3
NP_001036284.1
NP_649428.1
NP_476934.2
NP_001262566.1
NP_001261761.1
NP_001097153.1
NP_001285984.1
NP_651521.1
NP_650706.1
NP_608957.1
NP_651286.3
NP_001036264.2
NP_001261816.1
NP_611456.1
NP_001262064.1
NP_477445.1
NP_001259689.1
NP_001188578.1
NP_611264.2
NP_001287390.1
NP_001286308.1
NP_001262406.1
NP_649869.1
NP_648509.1
NP_609787.2
NP_649511.2
NP_001014641.2
NP_001096969.1
NP_001097501.1
NP_001261083.1
NP_001285680.1
NP_731744.1
NP_524938.1
NP_001138101.1
NP_648027.3
NP_001262533.1

| **Supplementary data 1C** | | |
|---|---|---|
| **Thermal adaptation genes specific to *Aedes koreicus*** | | |
| accession | uniprot accession | gene symbol |
| NP_001163722.1 | Q9VBX0 | Bili |
| NP_523578.1 | Q9NFV8 | Tep1 |
| NP_001027104.1 | Q0E8J1 | Svil |
| NP_001189037.1 | M9MRZ7 | Shab |
| NP_001097333.1 | P24014 | sli |
| NP_001246234.1 | A0A0B4K7L1 | Pfk |
| NP_001287456.1 | A0A0B4LHR4 | GABA-B-R2 |
| NP_001286665.1 | A0A0B4LG05 | tud |
| NP_001014535.1 | A1ZBH0 | Dmel_CG15117 |
| NP_648662.1 | Q9VU89 | Dmel_CG10710 |
| NP_001259851.1 | M9PAZ4 | IA-2 |
| NP_001097322.1 | Q1EC92 | Stacl |
| NP_001262456.1 | Q9VH09 | Dmel_CG3999 |

| **Supplementary data 1D** | | |
|---|---|---|
| **Thermal adaptation genes specific to *Aedes japonicus*** | | |
| accession | uniprot accession | gene symbol |
| NP_001260059.1 | F0JAH7 | hoe1 |
| NP_001262200.1 | P02574 | Act79B |
| NP_001188875.1 | A0A0B4JCQ5 | Lpin |
| NP_001096975.1 | Q9VY99 | Nna1 |
| NP_001260388.1 | Q9VKF6 | Vha100-5 |
| NP_001285589.1 | Q9VQR8 | LeuRS |
| NP_730938.1 | Q9VNC7 | NKCC |
| NP_609566.1 | Q9VK89 | CG6388 |
| NP_609040.1 | Q9VMC6 | 152078_at |
| NP_001263109.1 | Q9VA70 | Cdase |
| NP_723592.1 | Q9VKV0 | RluA-1 |
| NP_001261411.1 | M9PHD2 | Ctl1 |

| | | |
|---|---|---|
| NP_001260930.1 | Q7K0E6 | AspRS |
| NP_523364.2 | P54352 | eas |
| NP_001097369.2 | A8DYI4 | Sik3 |
| NP_001014621.1 | P20659 | trx |
| NP_001027070.1 | Q9VWX4 | CG16958 |
| NP_001014559.1 | Q9VZD1 | DopEcR |
| NP_001097556.2 | P48610 | Argk1 |
| NP_001262553.1 | Q8I0D4 | Dmel_CG9297 |
| NP_001262557.1 | Q95V55 | foxo |
| NP_001247239.1 | Q9VCY8 | AdipoR |
| NP_650086.2 | Q9VGM1 | Mrp4 |
| NP_649895.1 | Q9VHD2 | GstZ2 |
| NP_001286589.1 | Q8MRE7 | Hs3st-A |
| NP_001262131.1 | M9PG87 | CG3116 |
| NP_788515.1 | Q9VV61 | CG10162 |
| NP_001036284.1 | Q0KHR2 | f |
| NP_476934.2 | Q23979 | Myo61F |
| NP_001262566.1 | A0A0B4KFZ3 | Pde6 |
| NP_611456.1 | A1ZBR6 | Dme_CG11044 |
| NP_001262064.1 | Q9VW37 | SREBP |
| NP_001014641.2 | Q94900 | GluClalpha |
| NP_731744.1 | Q9VG26 | CG10135 |
| NP_001262533.1 | A0A0B4KFY8 | Dmel_CG9813 |

| **Supplementary data 1E** | | |
|---|---|---|
| **Thermal adaptation genes shared in *Aedes japonicus* and *Aedes koreicus*** | | |
| accession | uniprot accession | gene symbol |
| NP_001245529.1 | Q9W4K2 | CG3556 |
| NP_001286607.1 | A0A0B4LFX4 | cora |
| NP_001163731.1 | Q9VBP9 | Npl4 |
| NP_001262509.1 | Q2QBM1 | Men |
| NP_001245791.1 | Q04499 | slgA |
| NP_651004.1 | Q9VD53 | Dmel\CG13409 |
| NP_001260023.1 | Q7KU45 | l(2)k05819 |
| NP_728468.1 | Q9VR69 | Cda4 |
| NP_001027059.1 | Q9W3U8 | CG4564 |
| NP_001163749.1 | E1JIZ5 | Men-b |
| NP_001097890.1 | O77237 | Pli |
| NP_001284920.1 | X2JIJ6 | Grip |
| NP_610185.1 | Q8T0G1 | ZnT41F |
| NP_001036349.2 | Q0E8S1 | Dmel\CG32982 |
| NP_001097501.1 | A8JNK4 | Cip4 |
| NP_001259495.1 | Q9I7S6 | Pde9 |
| NP_001246206.1 | A0A0B4K715 | RyR |
| NP_523600.5 | P05031 | Ddc |
| NP_001262717.1 | Q8T390 | EndoA |
| NP_610724.1 | Q7K3N4 | Dmel\CG8888 |
| NP_001163667.1 | E1JIR4 | Atpalpha |
| NP_001261816.1 | Q7KUJ8 | Dmel\CG17364 |
