## Supplementary Table 2 for "De-novo genome assembly of the invasive mosquito species *Aedes japonicus* and *Aedes koreicus*"

Supplementary Data 2A: Enrichment analysis of 13 thermal adaptation genes specific to *Aedes koreicus*

kegg pathways

| Term | Overlap | P-value | Adjusted P-value | Genes |
|---|---|---|---|---|
| Pentose phosphate pathway | 1/24 | 0.015492787693 | 0.0370618720738194 | Pfk |
| Fructose and mannose metabolism | 1/29 | 0.018692433441 | 0.0370618720738194 | Pfk |
| Neuroactive ligand-receptor interaction | 1/54 | 0.034547169573 | 0.0370618720738194 | GABA-B-R2 |
| Galactose metabolism | 1/37 | 0.023791906863 | 0.0370618720738194 | Pfk |
| RNA degradation | 1/58 | 0.037061872073 | 0.0370618720738194 | Pfk |
| Glycolysis / Gluconeogenesis | 1/54 | 0.034547169573 | 0.0370618720738194 | Pfk |

Molecular Process

| Term | Overlap | P-value | Adjusted P-value | Genes |
|---|---|---|---|---|
| GABA receptor activity (GO:0016917) | 1/7 | 0.004541817095 | 0.0227090854774064 | GABA-B-R2 |
| voltage-gated potassium channel activity (GO:0005249) | 1/16 | 0.010353317603 | 0.0307087853313912 | Shab |
| voltage-gated cation channel activity (GO:0022843) | 1/19 | 0.012283514132 | 0.0307087853313912 | Shab |
| delayed rectifier potassium channel activity (GO:0005251) | 1/6 | 0.003894153996 | 0.0227090854774064 | Shab |
| protein tyrosine phosphatase activity (GO:0004725) | 1/28 | 0.018053273244 | 0.0354087263546230 | IA-2 |
| potassium channel activity (GO:0005267) | 1/33 | 0.021245235812 | 0.0354087263546230 | Shab |

Supplementary Data 2B: Enrichment analysis of 35 thermal adaptation genes specific to *Aedes japonicus*

Kegg pathways

| Term | Overlap | P-value | Adjusted P-value | Genes |
|---|---|---|---|---|
| Glycerophospholipid metabolism | 2/63 | 0.00543424724661338998804582800 | 0.0448273018166509620852 | eas;Lpin |
| Aminoacyl-tRNA biosynthesis | 2/64 | 0.0056034127270813702606511164 | 0.0448273018166509620852 | LeuRS;AspRS |

GO biological Process

| Term | Overlap | P-value | Adjusted P-value | Genes |
|---|---|---|---|---|
| cellular response to starvation (GO:0009267) | 4/80 | 0.0000113075145080456350167022 | 0.0028721086850435912501 | foxo;DopEcR;Sik3;Lpin |
| fatty acid metabolic process (GO:0006631) | 3/42 | 0.0000537840355859097195073345 | 0.0044944263143569520776 | AdipoR;SREBP;Lpin |
| lipid homeostasis (GO:0055088) | 3/40 | 0.0000463991406116026573660388 | 0.0044944263143569520776 | SREBP;Sik3;Lpin |
| cellular response to insulin stimulus (GO:0032869) | 3/46 | 0.0000707783671552275861190220 | 0.0044944263143569520776 | foxo;Sik3;Lpin |

Supplementary Data 2C: Enrichment analysis of 22 thermal adaptation genes in commun *Aedes japonicus* and *Aedes koreicus*

Kegg pathways

| Term | Overlap | P-value | Adjusted P-va | Genes |
|---|---|---|---|---|
| Tryptophan metabolism | 1/21 | 0.0228589677 | 0.0457179354 | Ddc |
| Phenylalanine metabolism | 1/8 | 0.0087677229 | 0.0350708919 | Ddc |
| Tyrosine metabolism | 1/21 | 0.0228589677 | 0.0457179354 | Ddc |
| Pyruvate metabolism | 2/46 | 0.0011609456 | 0.0092875652 | Men-b;Men |

Molecular function

| Term | Overlap | P-value | Adjusted P-va | Genes |
|---|---|---|---|---|
| malate dehydrogenase (decarboxylating) (NADP+) activity (GO:0004473) | 2/6 | 0.0000172797 | 0.0002505557 | Men-b;Men |
| malate dehydrogenase activity (GO:0016615) | 2/10 | 0.0000517010 | 0.0004997767 | Men-b;Men |
| malic enzyme activity (GO:0004470) | 2/6 | 0.0000172797 | 0.0002505557 | Men-b;Men |
| calcium-release channel activity (GO:0015278) | 1/7 | 0.0076757841 | 0.0259906299 | RyR |
| oxidoreductase activity, acting on the CH-OH group of donors, NAD or NADP as acceptor (GO:0016616) | 2/39 | 0.0008350521 | 0.0060541277 | Men-b;Men |
| lysophosphatidic acid acyltransferase activity (GO:0042171) | 1/7 | 0.0076757841 | 0.0259906299 | EndoA |
| glutamate receptor binding (GO:0035254) | 1/6 | 0.0065826972 | 0.0259906299 | Grip |
| lysophospholipid acyltransferase activity (GO:0071617) | 1/7 | 0.0076757841 | 0.0259906299 | EndoA |
| FAD binding (GO:0071949) | 1/10 | 0.0109481607 | 0.0264580552 | sIgA |
| 3',5'-cyclic-nucleotide phosphodiesterase activity (GO:0004114) | 1/9 | 0.0098585148 | 0.0259906299 | Pde9 |
| carboxy-lyase activity (GO:0016831) | 1/16 | 0.0174620379 | 0.0337599400 | Ddc |
| sodium:potassium-exchanging ATPase activity (GO:0005391) | 1/9 | 0.0098585148 | 0.0259906299 | Atpalpha |
| potassium-transporting ATPase activity (GO:0008556) | 1/9 | 0.0098585148 | 0.0259906299 | Atpalpha |
| divalent inorganic cation transmembrane transporter activity (GO:0072509) | 1/18 | 0.0196242154 | 0.0355688904 | ZnT41F |
| zinc ion transmembrane transporter activity (GO:0005385) | 1/16 | 0.0174620379 | 0.0337599400 | ZnT41F |
| ion transmembrane transporter activity (GO:0015075) | 1/14 | 0.0152953091 | 0.0337599400 | Atpalpha |
| flavin adenine dinucleotide binding (GO:0050660) | 1/29 | 0.0314352416 | 0.0479801056 | sIgA |
| transition metal ion transmembrane transporter activity (GO:0046915) | 1/22 | 0.0239349526 | 0.0408302133 | ZnT41F |
| ubiquitin binding (GO:0043130) | 1/26 | 0.0282275933 | 0.0454777892 | Npl4 |
