## Supplementary table 3 for "De-novo genome assembly of the invasive mosquito species *Aedes japonicus* and *Aedes koreicus*"

Supplementary data 3A: List of Insecticide resistance genes in *Aedes koreicus*

A0A6I8T7B5
A0A6I8TF11
Q174R4
A0A6I8TB15
A0A1S4FQ34
A0A1S4F2J6
Q16M87
A0A1S4G2F8
A0A1S4FQ35
Q17JR9
A0A6I8TJF9
A0A6I8TA62
A0A1S4FBX0
A0A1S4FQD9
A0A6I8TEM4
A0A6I8TKR2
A0A1S4EZP2
A0A6I8TBS9
A0A1S4F0H4
A0A6I8T699
A0A1S4FAP5
A0A1S4G2N2
A0A1S4FZY0
A0A6I8TA26
Q175A7
A0A6I8TPG0
A0A6I8T4K9
A0A1S4EXT0
Q16JY6
Q17DS9
A0A1S4F7G0
A0A6I8TG41
Q16XN3
A0A1S4FI93
Q17BQ1
A0A1S4FZV2
A0A1S4EX68
A0A1S4G5Q1
Q1HQV5
A0A6I8T781
A0A6I8TAK2
A0A6I8TJJ3
A0A6I8TGK2
A0A1S4FI55
Q16J86
A0A1S4G257
A0A6I8TR30
A0A6I8TPN3
Q17JE8

Q16RZ0
A0A1S4F3X1
Q16FX4
A0A6I8T7K3
Q16I47
Q16MY7
Q16P46
Q17G39
X5HYQ8
A0A6I8TQR9
Q16FX2
A0A6I8T9M3
A0A1S4FBC6
Q178T8
Q16L49
A0A6I8TNS7
A0A1S4FHK7
A0A1S4F7F6
A0A6I8T5K0
A0A1S4EWK0
A0A6I8TAD8
A0A1S4FKH1
A0A1S4F657
A0A6I8TGG4
Q16WR6
A0A1S4FVX1
Q16GD8
Q170T1
A0A1S4FHP0
Q17P07
Q177R6
A0A1S4FQK1
Q16WR5
A0A1S4FI63
A0A6I8TCE7
Q16F58
A0A1S4G2H7
A0A1S4G267
Q16QD6
A0A1S4FIQ7
Q16HC9
Q174X5
Q16HB6
A0A6I8TRE5
A0A1S4FLF9
A0A0P6IXI5
Q16QD7
Q178A6
A0A1S4G242
A0A1S4FBH9
A0A6I8T895
Q16WS1

Q173V4
A0A1S4F1N2
A0A1S4G0U5
A0A6I8TKI0
Q170T7
A0A1S4F3V0
A0A6I8TLX1
A0A1S4EWZ9
A0A1S4FUM7
Q170S9
Q17N26
A0A6I8TFB8
Q17GK7
Q16GE0
A0A1S4F6U9
A0A1S4FX07
Q17DY2
Q16XG2
A0A6I8TB20
A0A1S4FJZ4
A0A6I8TL04
A0A1S4F8B3
Q16FQ0
A0A6I8TKS0
A0A1S4EVV8
A0A1S4EX18
A0A1S4F3G0
A0A1S4FVG5
A0A6I8T9R3
Q16K89
A0A6I8TJP9
A0A1S4EW18
Q17JB0
A0A1S4FC72
A0A1S4F9A7
Q17KR8
A0A6I8TBS4
A0A1S4F313
A0A1S4EWV8
A0A6I8T9K3
Q17DN2
A0A1S4FLE3
A0A6I8TJH1
Q1HQF3
Q4U3X4
A0A1S4FK05
A0A1S4EZ49
A0A6I8TLW6
A0A6I8T3I5
Q17BK7
Q17GK4
Q16R02

A0A6I8TND4
Q179H3
Q16ZZ2
Q17DM8
Q16WR7
A0A1S4F8B2
Q176I8
A0A1S4EXM3
A0A6I8TJQ4
Q16ML8
A0A1S4EUN4
A0A6R5HP02
A0A1S4F6Y1
Q1HRC7
A0A1S4FWB3
Q16R00
Q1HRE3
Q16VA4
A0A6I8THT0
A0A6I8TE47
Q16XE6
A0A6I8T940
A0A1S4FNU3
Q17J69
Q176K1
Q16GD9
Q176J0
A0A6I8TCP5
Q176K5
Q176K0
A0A1S4FWJ8
Q16VA3
A0A6I8TCG0
Q17D91
Q1HQX7
A0A1S4FK40
A0A1S4F2K0
A0A6I8TKY9
A0A6I8TDN8
A0A6I8TAE0
A0A6I8T9Q6
A0A6I8T773
A0A6I8TKP9
A0A6I8T727
A0A1S4FSI4
A0A6I8T5Z0
A0A6I8T9A5
A0A1S4FCL9
A0A6I8TRP2
Q17IM5
A0A1S4F552
A0A1S4FL64

Q16EV5
Q176V4
A0A6I8TJX7
Q16WR9
Q16WS2
A0A1S4FUV2
Q16F48
Q16FA9
Q171F3
Q16ZI8
J9HHL7
A0A6I8T916
Q95W09
A0A6I8TBF7
A0A1S4FHX9
A0A1S4F244
A0A1S4FL91
Q17BT9
A0A1S4F753
A0A6I8TB50
Q16G28
A0A1S4FWY8
A0A1S4EVA2
A0A1S4F685
Q16TZ5
A0A1S4FRB0
A0A6I8T624
Q16J03
A0A1S4F5S1
A0A1S4EXS1
Q17DF6
A0A6I8TGW2
A0A1S4EUX1
A0A6I8TI39
A0A1S4F9K6
A0A1S4FHU5
A0A1S4EXP8
A0A1S4FG90
Q17D32
Q17PE2
Q16P41
A0A6I8TK63
A0A6I8TFC9
A0A6I8T9V7
A0A1S4FXL7
A0A6I8TJ74
A0A6I8TAA5
A0A6I8TDP5
A0A1S4EXN8
A0A6I8TKK8
A0A1S4FV60
Q17C03

A0A6I8T326
A0A6I8TK68
A0A1S4EXR7
A0A6I8TCS1
Q17P04
Q0IGD5
Q16IW1
Q16SH6
Q1DH17
A0A1S4FZG9
A0A6I8TE35
Q16X29
A0A6I8T3A2
A0A1S4G309
A0A1S4FJB1
A0A1S4FJ08
A0A1S4G2T5
A0A6I8TJW2
A0A6I8TH03
A0A1S4EX77
Q16W35
A0A1S4FJ19
Q16SJ1
Q16UT6
A0A1S4F3F3
A0A6I8T475
A0A6I8TDP1
A0A1S4FV52
A0A1S4FHI9
A0A1S4FPS6
A0A1S4FNB7
A0A1S4FIL0
Q17C00
Q174Q1
A0A6I8TAN6
A0A1S4FNC1
A0A6I8TLR0
A0A6I8TL99
Q16Z75
A0A1S4FVQ3
A0A1S4FQ43
Q17G14

Supplementary data 3B: List of Insecticide resistance genes in *Aedes japonicus*

A0A1S4EUN4
A0A1S4EUX1
A0A1S4EVV8
A0A1S4EW18
A0A1S4EWK0
A0A1S4EWV8
A0A1S4EWZ9
A0A1S4EX18
A0A1S4EX68
A0A1S4EX77
A0A1S4EXM3
A0A1S4EXN8
A0A1S4EXP8
A0A1S4EXR7
A0A1S4EXS1
A0A1S4EXT0
A0A1S4EYS0
A0A1S4EZ13
A0A1S4EZ49
A0A1S4EZV6
A0A1S4F0H4
A0A1S4F244
A0A1S4F2J6
A0A1S4F2K0
A0A1S4F313
A0A1S4F3F3
A0A1S4F3L2
A0A1S4F3V0
A0A1S4F3W2
A0A1S4F3W3
A0A1S4F4A4
A0A1S4F4H4
A0A1S4F552
A0A1S4F5S1
A0A1S4F5Y1
A0A1S4F657
A0A1S4F685
A0A1S4F6P9
A0A1S4F6U9
A0A1S4F6Y1
A0A1S4F753
A0A1S4F7F6
A0A1S4F7G0
A0A1S4F8A8
A0A1S4F8B2
A0A1S4F8B3
A0A1S4F9A7
A0A1S4F9D3
A0A1S4F9G0
A0A1S4F9K6
A0A1S4FAP5
A0A1S4FAQ9

A0A1S4FBC6
A0A1S4FBH9
A0A1S4FBX0
A0A1S4FC72
A0A1S4FCL9
A0A1S4FCV9
A0A1S4FEN8
A0A1S4FG90
A0A1S4FGV4
A0A1S4FHD7
A0A1S4FHK7
A0A1S4FHP0
A0A1S4FHU5
A0A1S4FHX9
A0A1S4FI63
A0A1S4FI93
A0A1S4FIB3
A0A1S4FIL0
A0A1S4FIQ7
A0A1S4FJB1
A0A1S4FJZ4
A0A1S4FK05
A0A1S4FK40
A0A1S4FKH1
A0A1S4FL58
A0A1S4FL64
A0A1S4FL91
A0A1S4FLF9
A0A1S4FLR3
A0A1S4FNC1
A0A1S4FNU3
A0A1S4FPF3
A0A1S4FPS6
A0A1S4FQ18
A0A1S4FQ34
A0A1S4FQ35
A0A1S4FQ39
A0A1S4FQD9
A0A1S4FQK1
A0A1S4FRH3
A0A1S4FSI4
A0A1S4FTE7
A0A1S4FU48
A0A1S4FUM7
A0A1S4FUR5
A0A1S4FUS1
A0A1S4FUV2
A0A1S4FV52
A0A1S4FV60
A0A1S4FVQ3
A0A1S4FVW1
A0A1S4FVX1

A0A1S4FW30
A0A1S4FWB3
A0A1S4FWJ8
A0A1S4FWU1
A0A1S4FWY8
A0A1S4FX07
A0A1S4FXL7
A0A1S4FZB8
A0A1S4FZE2
A0A1S4FZE7
A0A1S4FZG9
A0A1S4FZV2
A0A1S4FZY0
A0A1S4G0U5
A0A1S4G252
A0A1S4G257
A0A1S4G267
A0A1S4G2B1
A0A1S4G2C1
A0A1S4G2F8
A0A1S4G2N2
A0A1S4G2T5
A0A1S4G560
A0A1S4G5Q1
A0A6I8T340
A0A6I8T3B0
A0A6I8T3H9
A0A6I8T3I5
A0A6I8T475
A0A6I8T4K9
A0A6I8T4R9
A0A6I8T5J8
A0A6I8T5K0
A0A6I8T5X6
A0A6I8T5Z0
A0A6I8T605
A0A6I8T624
A0A6I8T699
A0A6I8T6U0
A0A6I8T727
A0A6I8T773
A0A6I8T781
A0A6I8T7D2
A0A6I8T7K3
A0A6I8T7Q9
A0A6I8T836
A0A6I8T861
A0A6I8T865
A0A6I8T895
A0A6I8T8T8
A0A6I8T940
A0A6I8T9A5

A0A6I8T9K3
A0A6I8T9N7
A0A6I8T9Q6
A0A6I8T9R3
A0A6I8T9V7
A0A6I8TA26
A0A6I8TA31
A0A6I8TA43
A0A6I8TA62
A0A6I8TAA5
A0A6I8TAD8
A0A6I8TAE0
A0A6I8TAK2
A0A6I8TAK3
A0A6I8TAN1
A0A6I8TAN6
A0A6I8TB20
A0A6I8TB50
A0A6I8TBF7
A0A6I8TBS3
A0A6I8TBS4
A0A6I8TC40
A0A6I8TCE7
A0A6I8TCS1
A0A6I8TCY2
A0A6I8TD77
A0A6I8TDL8
A0A6I8TDN8
A0A6I8TDP1
A0A6I8TDP3
A0A6I8TE12
A0A6I8TE47
A0A6I8TE68
A0A6I8TEM4
A0A6I8TET9
A0A6I8TEU9
A0A6I8TF13
A0A6I8TF78
A0A6I8TFB8
A0A6I8TFC9
A0A6I8TFJ6
A0A6I8TFL6
A0A6I8TG41
A0A6I8TGK2
A0A6I8TGW2
A0A6I8THT0
A0A6I8THW7
A0A6I8TI39
A0A6I8TJ74
A0A6I8TJA0
A0A6I8TJC5
A0A6I8TJF9

A0A6I8TJH1
A0A6I8TJJ3
A0A6I8TJM9
A0A6I8TJX7
A0A6I8TK63
A0A6I8TK68
A0A6I8TKI0
A0A6I8TKP1
A0A6I8TKP8
A0A6I8TKP9
A0A6I8TKR2
A0A6I8TKS0
A0A6I8TKY9
A0A6I8TL04
A0A6I8TL99
A0A6I8TLA6
A0A6I8TLK7
A0A6I8TLR0
A0A6I8TLW6
A0A6I8TLX1
A0A6I8TNS7
A0A6I8TNZ7
A0A6I8TPG0
A0A6I8TPN3
A0A6I8TQR9
A0A6I8TRP2
A0A6R5HP02
J9HGP0
Q0IG96
Q0IG97
Q0IG98
Q0IGD5
Q16EV5
Q16F48
Q16FA9
Q16FQ0
Q16FX2
Q16FX4
Q16G28
Q16GD8
Q16GD9
Q16GE0
Q16HB6
Q16HC9
Q16HI0
Q16I47
Q16IW1
Q16J02
Q16J03
Q16J86
Q16JB8
Q16JY6

Q16K89
Q16L49
Q16M87
Q16ML8
Q16MY7
Q16P41
Q16P46
Q16P53
Q16QD6
Q16QD7
Q16QL8
Q16R00
Q16R02
Q16RE9
Q16RZ0
Q16SH6
Q16SJ1
Q16SL2
Q16T85
Q16UT5
Q16V50
Q16VA3
Q16VA4
Q16W35
Q16WQ4
Q16WR5
Q16WR6
Q16XE6
Q16XN3
Q16XV0
Q16Z75
Q16ZD7
Q16ZI8
Q16ZZ2
Q170C9
Q170S9
Q170T1
Q170T7
Q171F3
Q173V3
Q173V4
Q173V5
Q174Q1
Q174T2
Q174T4
Q176I8
Q176J0
Q176K0
Q176K1
Q176K6
Q176V4
Q177R6

Q177U0
Q178A6
Q178T8
Q179H3
Q17BE2
Q17BK7
Q17BQ1
Q17C00
Q17C04
Q17C17
Q17C20
Q17D32
Q17D91
Q17DF6
Q17DM8
Q17DM9
Q17DN2
Q17DS9
Q17DY2
Q17G14
Q17G39
Q17G40
Q17GK4
Q17GK7
Q17HK2
Q17IM5
Q17J69
Q17JB0
Q17JE8
Q17JN6
Q17K53
Q17KM5
Q17KR8
Q17KZ6
Q17N26
Q17P04
Q17P07
Q17PE2
Q1DH17
Q1HQX7
Q1HRE3
Q45KG6
Q4U3X4
Q5K6H6
Q5PY78
Q95W09
Q9GYW1
X5HYQ8

Supplementary data 3C: Insecticide resistance genes specific to *Aedes koreicus*

A0A6I8TEM4
A0A1S4G2N2
A0A1S4F7G0
A0A6I8T781
A0A1S4FBC6
A0A6I8TLX1
A0A1S4F9A7
Q17BK7
Q179H3
A0A6I8TK63
A0A6I8TDP1
Q17C00
A0A1S4FVQ3

Supplementary data 3D: Insecticide resistance genes specific to *Aedes japonicus*

A0A6I8TA43
A0A1S4F5Y1
A0A6I8T340
A0A1S4FVW1
A0A6I8T5X6
A0A6I8TET9
Q173V5
A0A1S4FRH3
A0A6I8T5J8
Q17C04
