## Supplementary Table 4 for "De-novo genome assembly of the invasive mosquito species *Aedes japonicus* and *Aedes koreicus*"

Supplementary data 4A – Differentially expressed transcripts in adults of *Aedes japonicus*

| ID | log2FoldChange | pvalue | padj | Gene symbol |
|---|---|---|---|---|
| NODE_20766_length_2483_cov_163.774274_g6364_i3 | 9.99847753374144 | 0.000485812869457969 | 0.0444217333247133 | AEF1_DROME |
| NODE_90229_length_503_cov_30708.076744_g52874_i0 | 9.87244632560482 | 0.000268823470748356 | 0.0394189551753458 | 5HTB2_APLCA |
| NODE_7499_length_4262_cov_613.735498_g2098_i5 | 9.86883078027984 | 0.000638442482752707 | 0.048387292635351 | ACSL6_MOUSE |
| NODE_8896_length_3950_cov_129.869745_g2319_i2 | 9.79868839249711 | 0.000513245058641282 | 0.0451938212669208 | AAP1_YEAST |
| NODE_42290_length_1353_cov_332.074219_g14637_i0 | 9.74002122627872 | 0.000699807265607805 | 0.0487954959510264 | A2GL_HUMAN |
| NODE_13298_length_3228_cov_288.749604_g337_i32 | 9.70123399647226 | 0.000752347867427485 | 0.0491500250048068 | AEF1_DROME |
| NODE_59394_length_871_cov_225.204261_g24843_i0 | 9.67268195277328 | 0.000822347558658119 | 0.0497407192834807 | ACRO_MOUSE |
| NODE_33578_length_1718_cov_267.552584_g8952_i1 | 9.56093355747999 | 0.00074527880813417 | 0.0489855485219545 | ACBP4_ARATH |
| NODE_44855_length_1263_cov_541.475630_g5383_i4 | 9.53032002361476 | 0.000343173205248018 | 0.0410216895935492 | ACET6_CERPU |
| NODE_5455_length_4824_cov_71.041675_g2134_i0 | 9.53012289269309 | 0.000721003267393825 | 0.0489855485219545 | 172L_IIV6 |
| NODE_44229_length_1284_cov_466.733278_g15524_i4 | 9.49749194958119 | 0.000574353721444508 | 0.046446184647473 | ACRO_HUMAN |
| NODE_44984_length_1259_cov_1567.225126_g15908_i0 | 9.4853729518457 | 0.000336198009450157 | 0.0410216895935492 | ACON_CAEEL |
| NODE_10618_length_3626_cov_1768.617506_g3293_i2 | 9.34321975885047 | 0.000452094083300618 | 0.0438115605171869 | AFG2_SCHPO |
| NODE_31396_length_1821_cov_11856.094394_g10348_i0 | 9.03932266057271 | 0.000135981182643718 | 0.03229017112988 | ADT2_ANOGA |
| NODE_16871_length_2830_cov_408.499093_g3943_i2 | 8.96947009587114 | 0.000563494970906171 | 0.0462845914228046 | AEF1_DROME |
| NODE_28960_length_1945_cov_64487.162927_g5743_i19 | 8.84952866444679 | 0.00045692143123123 | 0.0438115605171869 | ACT1_AEDAE |
| NODE_93816_length_482_cov_47890.650367_g56423_i0 | 8.84105030847384 | 0.000770872024493983 | 0.0493956502037889 | ACRO_HUMAN |
| NODE_34695_length_1671_cov_974.259074_g4313_i7 | 8.72766090083167 | 0.00040595647810079 | 0.0423394429503389 | ACRO_RABIT |
| NODE_31764_length_1803_cov_6371.989017_g10487_i0 | 8.1392817642137 | 0.000723914498248925 | 0.0489855485219545 | ACTN_ACTDE |
| NODE_77274_length_606_cov_104793.536585_g40243_i0 | 26.8258435707979 | 2.32830854558929e-14 | 5.95979828937645e-11 | 5HT7R_XENLA |
| NODE_20366_length_2515_cov_4063.611384_g3850_i11 | 13.6906131583633 | 9.14631410665135e-06 | 0.00840955443611294 | ABAQ_ACIBT |
| NODE_53229_length_1012_cov_60909.587859_g20591_i0 | 13.6005408279102 | 3.19395859210934e-06 | 0.00422352570197876 | A16L1_HUMAN |
| NODE_25535_length_2141_cov_82847.455513_g8353_i0 | 13.5766167416038 | 8.3522004075666e-05 | 0.0282769371248874 | ADT1_ANOGA |
| NODE_21042_length_2460_cov_2883.450775_g6910_i1 | 13.2662408095243 | 0.000119894597655766 | 0.0319740371131077 | 3O1D_MYCS2 |
| NODE_33175_length_1737_cov_11084.894231_g313_i160 | 13.0559954428957 | 0.000147925472715489 | 0.03229017112988 | ABRU_DROME |
| NODE_34827_length_1665_cov_11704.750000_g5417_i3 | 13.0543240477665 | 0.000146210181951804 | 0.03229017112988 | ACTNA_DICDI |
| NODE_54036_length_991_cov_87125.010893_g21117_i0 | 12.9938547278361 | 8.11247431322264e-05 | 0.0282769371248874 | ADRB2_TSCTR |
| NODE_33630_length_1716_cov_1853.273281_g2353_i17 | 12.9681225083967 | 6.75857017779383e-05 | 0.0270619250169047 | ACTNA_DICDI |
| NODE_58096_length_897_cov_1589.026699_g21899_i1 | 12.6543756342465 | 0.000227253436497313 | 0.0375023215450296 | ACRO_MELGA |
| NODE_32792_length_1756_cov_28185.212715_g10863_i0 | 12.6341569067083 | 1.96961635532817e-06 | 0.00306585915354453 | ACRO_HUMAN |
| NODE_27742_length_2011_cov_58341.738906_g5743_i14 | 12.5272024585374 | 0.000255002907369424 | 0.0389063169457655 | ACT1_AEDAE |
| NODE_62678_length_807_cov_390.861035_g27317_i0 | 12.3677936528891 | 0.000350058282762391 | 0.0410216895935492 | ACRO_HUMAN |
| NODE_37534_length_1543_cov_5607.436054_g4769_i3 | 12.342005487546 | 0.000307572163277138 | 0.0409947280184489 | A9A1A_DANRE |
| NODE_26017_length_2109_cov_1501.785363_g1701_i9 | 12.2634325573305 | 0.000333741588375964 | 0.0410216895935492 | ACTN_ACTDE |
| NODE_41075_length_1400_cov_3127.915599_g14102_i0 | 12.2575949080062 | 0.000336650616387133 | 0.0410216895935492 | ADAR_DROME |
| NODE_54429_length_981_cov_767.046256_g19900_i1 | 12.2551061055598 | 9.57214815850618e-05 | 0.0295965738504467 | AGLA_RHIME |
| NODE_48762_length_1139_cov_4670.465291_g17931_i0 | 12.2192801411173 | 4.42936277362383e-05 | 0.0211996583489922 | ACP1_ENTHI |
| NODE_60055_length_858_cov_778.020382_g25327_i0 | 12.1297903736677 | 0.000381493123044177 | 0.0419150573145927 | ACOB_BACSU |
| NODE_7755_length_4202_cov_15594.609833_g2328_i5 | 12.1189513004525 | 0.000386567959508012 | 0.0419188104699125 | AAAH_CHLPN |
| NODE_25111_length_2167_cov_4408.129417_g5031_i13 | 12.103630830161 | 7.12785051863052e-05 | 0.0273670295901573 | 332L_IIV6 |
| NODE_26699_length_2069_cov_206.657816_g8766_i0 | 12.0817140209222 | 0.000400313320421068 | 0.0423394429503389 | A2GL_HUMAN |
| NODE_27681_length_2015_cov_209.828012_g9074_i0 | 11.9700354073114 | 0.000454300817843841 | 0.0438115605171869 | 72A61_PARPY |
| NODE_59480_length_869_cov_450.157035_g24905_i0 | 11.9570659049337 | 0.00046717663080077 | 0.0441153629825625 | A1AT_MERUN |
| NODE_39634_length_1458_cov_2104.151625_g7955_i6 | 11.9455814514156 | 0.000462548354967422 | 0.0438115605171869 | ADH1_GEOSE |
| NODE_31667_length_1808_cov_1475.361960_g6352_i5 | 11.8628278301073 | 0.000511752594497974 | 0.0451938212669208 | 72A61_TRIFG |
| NODE_28749_length_1957_cov_57831.015924_g4496_i10 | 11.8533370757051 | 0.000510663499555702 | 0.0451938212669208 | ACT5C_ANOGA |
| NODE_33142_length_1739_cov_24734.250300_g10863_i1 | 11.8460787782395 | 1.71846978752603e-05 | 0.0117728658639945 | ACRO_RAT |
| NODE_34386_length_1684_cov_18528.399131_g5014_i9 | 11.8274911802341 | 0.000518072758222272 | 0.0452523602488199 | ABF2_YEAST |
| NODE_20465_length_2508_cov_16277.476797_g2432_i14 | 11.7998515716149 | 0.000607242376880428 | 0.0471891701207158 | A1CF_HUMAN |
| NODE_33243_length_1734_cov_11104.852498_g313_i161 | 11.7600586770914 | 0.000561079216314384 | 0.0462845914228046 | ABRU_DROME |
| NODE_35809_length_1620_cov_2070.248869_g7955_i4 | 11.7594952609824 | 0.000558791558512153 | 0.0462845914228046 | ADH1_GEOSE |
| NODE_5108_length_4934_cov_1965.814853_g1979_i3 | 11.7269588058355 | 0.000571068726088071 | 0.046381740843871 | AATM_MOUSE |
| NODE_67460_length_728_cov_411.035115_g31261_i0 | 11.6811025129879 | 0.000223706481713363 | 0.0374698088820983 | ACRO_SHEEP |
| NODE_11433_length_3490_cov_1344.093942_g1543_i3 | 11.6621494613786 | 3.18442501391832e-05 | 0.0176604167557606 | AFAD_MOUSE |
| NODE_13327_length_3225_cov_1760.598985_g1150_i10 | 11.6000778818547 | 0.000647633402706275 | 0.0484625319590834 | AGAP2_HUMAN |
| NODE_2300_length_6357_cov_457.193189_g988_i2 | 11.5945012976583 | 0.000675001131483865 | 0.0487954959510264 | ACBD6_BOVIN |
| NODE_15564_length_2964_cov_2168.245244_g5089_i3 | 11.5798164882635 | 3.97536442274944e-05 | 0.0198421796523489 | ACBP5_ARATH |
| NODE_39981_length_1443_cov_214.097080_g13612_i0 | 11.5678123671537 | 0.000723115660269343 | 0.0489855485219545 | ACRO_RABIT |
| NODE_46710_length_1204_cov_1759.679045_g15654_i1 | 11.5532241075029 | 0.000754011660396018 | 0.0491500250048068 | ACRO_MELGA |
| NODE_44671_length_1269_cov_510.433110_g15749_i0 | 11.5350675470907 | 0.000701998853117769 | 0.0487954959510264 | ACRO_MOUSE |
| NODE_6441_length_4537_cov_1249.943324_g2401_i1 | 11.5113968539413 | 0.00070763242206495 | 0.0487954959510264 | AFG2_YEAST |
| NODE_64231_length_780_cov_918.848656_g28569_i0 | 11.4656542111623 | 0.000739351365680739 | 0.0489855485219545 | ACTN_ACTDE |
| NODE_7798_length_4191_cov_439.539825_g2221_i4 | 11.4570972410354 | 5.50319042403945e-05 | 0.0252994697665151 | 095R_FRG3G |
| NODE_26948_length_2056_cov_554.171962_g8849_i0 | 11.4452377734414 | 0.000757294960026438 | 0.0491869157721051 | A2GL_HUMAN |
| NODE_26896_length_2059_cov_174.362538_g8827_i0 | 11.4250570154077 | 0.00082760147293547 | 0.0497686193853569 | 5NTD_DIPOM |
| NODE_892_length_8208_cov_2113.155993_g399_i0 | 11.4248930468104 | 0.000809641891449247 | 0.0497045888057081 | 13_ARATH, 5FCLL_AR |
| NODE_33178_length_1737_cov_1657.323918_g10540_i4 | 11.4020287153667 | 0.000388757960573116 | 0.0419188104699125 | ACRO_HUMAN |
| NODE_946_length_8104_cov_826.452995_g349_i5 | 11.3864305878883 | 5.07926954950389e-05 | 0.0236619465053489 | ACAC_CHICK |
| NODE_20753_length_2484_cov_612.608876_g3098_i10 | 11.3535407857673 | 0.000283826935353787 | 0.0394189551753458 | ADD_SHESW |
| NODE_30219_length_1880_cov_1914.938572_g5031_i14 | 11.2366753391476 | 0.000462269037449162 | 0.0438115605171869 | 095L_IIV6 |
| NODE_29108_length_1938_cov_1386.987131_g5697_i7 | 11.1860990073983 | 6.75361477643696e-05 | 0.0270619250169047 | ACTZ_DROME |
| NODE_20115_length_2536_cov_803.392205_g6459_i1 | 11.0668067532804 | 8.6826538270712e-05 | 0.0286191737796265 | AFG32_HUMAN |
| NODE_28517_length_1970_cov_1264.396415_g4875_i8 | 10.9887718458598 | 0.000427176478856123 | 0.0436113436451875 | AAAH_CHLPN |
| NODE_4513_length_5164_cov_1053.305441_g1582_i2 | 10.9762283908218 | 0.000367477089743873 | 0.0413052343751579 | ACRO_MOUSE |
| NODE_35061_length_1654_cov_1091.637571_g11169_i1 | 10.9729955027638 | 0.000145767144164378 | 0.03229017112988 | ACRO_RAT |
| NODE_1760_length_6860_cov_475.307352_g341_i3 | 10.8678722872569 | 0.000117031556925869 | 0.0319740371131077 | 7LESS_DROVI |
| NODE_32119_length_1787_cov_579.768961_g1852_i5 | 10.8133029414729 | 0.000140578843663209 | 0.03229017112988 | ACTNA_DICDI |
| NODE_21987_length_2382_cov_1211.074924_g6196_i5 | 10.8010424067115 | 0.000276648055554997 | 0.0394189551753458 | AFG2_SCHPO |
| NODE_19407_length_2596_cov_191.218391_g4981_i3 | 10.7931255975123 | 0.000119043393811966 | 0.0319740371131077 | ABPB_ECOLI |
| NODE_65256_length_763_cov_282.262319_g29418_i0 | 10.7737521741942 | 0.000535214941642237 | 0.0458403784586327 | ACBD6_DANRE |
| NODE_39814_length_1450_cov_8930.260712_g5743_i21 | 10.6969770057648 | 0.000430224024882972 | 0.0436113436451875 | ACT1_AEDAE |
| NODE_61278_length_833_cov_84715.265789_g26247_i0 | 10.6824463865382 | 0.000237519099815185 | 0.0378998207889699 | 5HT7R_XENLA |
| NODE_84006_length_546_cov_559.186047_g46757_i0 | 10.6289224722253 | 0.00053594389822598 | 0.0458403784586327 | ACRO_HUMAN |
| NODE_75236_length_627_cov_101416.019856_g38311_i0 | 10.6180523038909 | 0.000402713987874674 | 0.0423394429503389 | 5HT7R_XENLA |
| NODE_37517_length_1544_cov_1683.450714_g12592_i1 | 10.5328064392679 | 0.000607161499454094 | 0.0471891701207158 | AFG2_SCHPO |
| NODE_28929_length_1947_cov_140.459979_g9481_i0 | 10.4657985996037 | 0.000339233184662208 | 0.0410216895935492 | A2GL_HUMAN |
| NODE_27970_length_1999_cov_380.909657_g3312_i2 | 10.4228098281761 | 0.000206390446599701 | 0.0362365618781254 | 235L_IIV6 |
| NODE_17909_length_2732_cov_4553.696502_g5031_i3 | 10.3830054985142 | 0.000214496538877668 | 0.0367367380972884 | 332L_IIV6 |
| NODE_13651_length_3183_cov_229.001929_g2895_i2 | 10.3814706746277 | 0.000229295087227574 | 0.0376119298246206 | A16L1_HUMAN |
| NODE_17715_length_2751_cov_27.635922_g5896_i0 | 10.3796144758743 | 0.000630710336964043 | 0.0483254132964621 | AAP1_YEAST |
| NODE_5912_length_4675_cov_289.379835_g1880_i1 | 10.3754373835829 | 0.000234067403755541 | 0.0378998207889699 | AEF1_DROME |
| NODE_28328_length_1980_cov_1538.680650_g9056_i1 | 10.3056255390135 | 0.000705437982176516 | 0.0487954959510264 | 5GT_VERHY |
| NODE_20066_length_2541_cov_165.508104_g1212_i6 | 10.2921798555316 | 0.000682498875457625 | 0.0487954959510264 | ABR_XENTR |
| NODE_24498_length_2206_cov_2897.287389_g4847_i6 | 10.2317128812558 | 0.000192490518278736 | 0.0348467679696412 | AAT_PYRAB |
| NODE_14797_length_3055_cov_520.249832_g4590_i3 | 10.1811294860612 | 0.000489681075485529 | 0.0444217333247133 | AAP1_YEAST |
| NODE_33717_length_1712_cov_1247.387431_g11208_i0 | 10.1626773375278 | 0.000309757239671156 | 0.0409947280184489 | 3AHDP_RUMGV |
| NODE_32856_length_1753_cov_682.191667_g10651_i1 | 10.1540076631242 | 0.000341587700721053 | 0.0410216895935492 | AGAP1_HUMAN |
| NODE_4929_length_5005_cov_116.146796_g1581_i3 | 10.1345298816112 | 0.000313749703171385 | 0.0410216895935492 | AES_XENLA |
| NODE_10524_length_3641_cov_79.133688_g1568_i3 | 10.1128892062205 | 0.000663597646606083 | 0.0486933408602592 | A1CF_HUMAN |
| NODE_4420_length_5192_cov_302.855441_g1398_i4 | 10.0676677561338 | 0.000376554261925619 | 0.0416342701184152 | AB10C_ARATH |
| NODE_11241_length_3521_cov_867.041763_g719_i21 | 10.0580941774942 | 0.000607777170334643 | 0.0471891701207158 | AFC2_ARATH |
| NODE_2261_length_6388_cov_1358.008076_g601_i3 | 10.0429440367193 | 0.000508211454109785 | 0.0451938212669208 | ACES_BOMIG |
| NODE_20707_length_2488_cov_372.286128_g4753_i2 | 10.0256119709437 | 0.000355039081061146 | 0.0410216895935492 | A20A5_HUMAN |
| NODE_17568_length_2763_cov_2226.975093_g3463_i4 | -9.96936050430785 | 0.00040783281202817 | 0.0424085435102745 | ADNT1_ARATH |
| NODE_36373_length_1595_cov_954.709593_g7759_i3 | -9.85923686916147 | 0.000591234247992634 | 0.0469480304332151 | ACR1_ACIAD |
| NODE_6347_length_4567_cov_14936.168669_g2328_i3 | -9.85648907195387 | 0.000513522053950243 | 0.0451938212669208 | AAAH_CHLPN |
| NODE_17730_length_2749_cov_1338.848281_g4875_i6 | -9.81213113841882 | 0.000239728108828597 | 0.0378998207889699 | AAAH_CHLPN |
| NODE_30806_length_1851_cov_13700.683915_g313_i141 | -9.67823624205761 | 0.000601306359812252 | 0.047115856284359 | ABRU_DROME |
| NODE_25840_length_2122_cov_74.782333_g8468_i0 | -9.65601576740754 | 0.000601384332415994 | 0.047115856284359 | AEF1_DROME |
| NODE_28401_length_1977_cov_478.219538_g9142_i1 | -9.63850530879621 | 0.000648930961506894 | 0.0484625319590834 | AEF1_DROME |
| NODE_1705_length_6909_cov_323.028525_g742_i1 | -9.63160973447623 | 0.000745763339164705 | 0.0489855485219545 | AAPK1_CAEEL |
| NODE_12537_length_3330_cov_123.304882_g2811_i7 | -9.62310357234075 | 0.000636609722496134 | 0.048387292635351 | ACACA_SHEEP |
| NODE_2160_length_6480_cov_356.436398_g938_i0 | -9.61730541195022 | 0.000700164630492115 | 0.0487954959510264 | ACTNB_DICDI |
| NODE_16312_length_2886_cov_605.939211_g5474_i1 | -9.56108087602338 | 0.000684932348791275 | 0.0487954959510264 | AEF1_DROME |
| NODE_22863_length_2318_cov_124.890869_g7468_i0 | -9.50583424820012 | 0.000766921761185464 | 0.0493956502037889 | AEF1_DROME |
| NODE_8328_length_4068_cov_557.273842_g2980_i3 | -12.3605105026859 | 1.11321887428824e-05 | 0.0090452916857574 | AGC1_YEAST |
| NODE_11302_length_3512_cov_2375.917418_g2030_i6 | -11.7708352381972 | 2.68537531369093e-05 | 0.0159024284889911 | A2ML1_HUMAN |
| NODE_2139_length_6494_cov_2027.105591_g227_i15 | -11.5418909231934 | 0.000586351877917701 | 0.0467983398795417 | ACRS2_ALTAL |
| NODE_788_length_8467_cov_790.582321_g349_i1 | -11.2586154120088 | 7.70209587643993e-05 | 0.0278128854777649 | ACAC_CHICK |
| NODE_41901_length_1367_cov_2459.803709_g1158_i9 | -11.2376683643611 | 0.00082649937834292 | 0.0497686193853569 | A1CF_MOUSE |
| NODE_3362_length_5701_cov_1024.346304_g1371_i2 | -11.1594755392676 | 9.46502717883643e-05 | 0.0295965738504467 | ADD1_CAEEL |
| NODE_8942_length_3939_cov_1412.866529_g2401_i2 | -11.0585266788698 | 8.36776330734749e-05 | 0.0282769371248874 | AFG2_YEAST |
| NODE_4182_length_5295_cov_600.040597_g164_i4 | -10.9128475238632 | 0.000111963063556357 | 0.0319740371131077 | ABR_MOUSE |
| NODE_25895_length_2117_cov_9398.594912_g8486_i0 | -10.7323406206636 | 0.000142513676872094 | 0.03229017112988 | ACRO_RAT |
| NODE_9734_length_3786_cov_797.315378_g3517_i0 | -10.5890054397152 | 0.000173270423185691 | 0.0327065987215749 | CF1_PIG, A1CF_MOU |
| NODE_6506_length_4518_cov_613.391676_g2098_i3 | -10.4882625662307 | 0.000251461869447495 | 0.0387040804256653 | ACSL6_MOUSE |
| NODE_4183_length_5295_cov_560.588280_g1044_i4 | -10.3798579730701 | 0.000220806520587819 | 0.0374502865185331 | A1CF_RAT |
| NODE_20558_length_2500_cov_434.868150_g612_i3 | -10.2918037324147 | 0.00030298690581027 | 0.0409947280184489 | AEF1_DROME |
| NODE_20802_length_2480_cov_380.638970_g3312_i1 | -10.1635215810005 | 0.000358790857801876 | 0.0411009632155401 | 235L_IIV6 |
| NODE_898_length_8199_cov_212.357741_g401_i0 | -10.1574019657631 | 0.000364455071360393 | 0.0413052343751579 | A1CF_MOUSE |
| NODE_1118_length_7783_cov_322.580285_g497_i0 | -10.0954451171097 | 0.000339195730795317 | 0.0410216895935492 | ACA7_ORYSJ |
| NODE_10301_length_3686_cov_284.912538_g3695_i0 | -10.0503373437979 | 0.000371584673992789 | 0.0413052343751579 | 6PGL_RAT |
| NODE_691_length_8725_cov_388.824896_g304_i0 | -10.0323481724473 | 0.000356924744166661 | 0.0410216895935492 | ADT2_CAEEL |

Supplementary data 4B – Differentially expressed transcripts in larvae of *Aedes japonicus*

| ID | log2FoldChange | pvalue | padj | Gene symbol |
|---|---|---|---|---|
| NODE_1_length_524078_cov_28.869238_g0_i0 | 12.7231338904642 | 4.23976618804311e-13 | 4.09004187351679e-09 | ADDB_STRA3,ADIPO_MOUSE,AFSQ2_STRCO,ACP_NOVAD,ACOR_BACSU,AGL11_HALVD,AGLB_THENE,ABAF_ACIBT,AGLA_RHIME,AAPA_BACSU,ACOR_BACSU |
| NODE_10_length_97030_cov_28.306486_g5_i0 | 12.4873737976679 | 9.6638109760878e-12 | 2.50991472151253e-08 | ACRB_ECOLI,ACCA1_MYCTU,ABCBA_HUMAN,AEPA_PECCC,ACCD1_MYCTU,ABDH_PECAS,ACDA_BACSU,AGL9_HALVD,ACRA_ECOLI,ACNA_BRADU,ACSE_MOO |
| NODE_10062_length_3732_cov_13.710577_g3621_i0 | 9.0087548942671 | 6.15436609599584e-06 | 0.00171732245343143 | AFG2_SCHPO,ABAB_STRAT |
| NODE_101_length_16277_cov_12.784991_g34_i0 | 10.0325480795006 | 7.54934616438954e-08 | 4.67699309898071e-05 | ACDA_METBU |
| NODE_1017_length_7971_cov_10.977336_g451_i0 | 10.007941224912 | 6.15453581848841e-07 | 0.000280170268433522 | AGL9_HALVD,AARE1_ORYSJ |
| NODE_10894_length_3581_cov_14.198119_g3859_i0 | 9.05953900335962 | 5.13623643164141e-06 | 0.00148858271998232 | AGLUS_THESQ |
| NODE_11_length_89169_cov_27.482435_g6_i0 | 12.2496757970894 | 5.2244147445411e-13 | 4.40992848586715e-09 | ABCBA_MOUSE,AFSQ2_STRCO,ACOX4_ARATH,ACRE_ECOLI,ACRE_ECOLI,ACRD_ECOLI,ACOCT_ACEAC,AAEA_PECAS,ACRD_ECOLI,ABGR_ECOLI,AFSQ1_STR |
| NODE_110_length_15335_cov_20.721531_g36_i0 | 5.61336102762534 | 2.08896522368244e-05 | 0.00486426357326992 | ABR1_ASPFU,ADH_MORSE,ACLDH_PSEAI |
| NODE_12_length_86351_cov_30.594960_g4_i4 | 11.27727150434 | 1.5817820539424e-10 | 2.60523362289324e-07 | 4CLL7_ARATH,AFSQ1_STRCO,2OCS_HYDTT,ACRT_ASPA1,ADRX_YEAST,AAEA_ERWT9,ADCS_STRVP,ACOR_BACSU,ADCL2_HUMAN,AGAA_GEOSE,ABD12_RAT |
| NODE_121_length_14686_cov_13.019366_g41_i0 | 9.58276353678092 | 3.05978683036724e-07 | 0.000151288737068256 | AFR_ENSAD,ADHA_MYCTU |
| NODE_1240_length_7589_cov_12.291911_g550_i0 | 9.92801505786081 | 8.76895788975522e-07 | 0.000365524807641599 | AFSQ2_STRCO,ACEA_KOMXY,ACOR_BACSU |
| NODE_12576_length_3325_cov_1583.381919_g956_i4 | -8.85788720034276 | 1.54595380463569e-07 | 8.69959737661993e-05 | AGAP2_HUMAN |
| NODE_12643_length_3319_cov_11.921134_g4379_i0 | 8.67106285862709 | 1.44554902210873e-05 | 0.00356259249507147 | ABCH2_DICDI |
| NODE_1271_length_7544_cov_11.840851_g561_i0 | 10.0202628410485 | 6.66738211076971e-07 | 0.000296052885894764 | AAER_ECOLI |
| NODE_12870_length_3287_cov_9.362166_g4454_i0 | 9.68165702355873 | 1.07302146760116e-06 | 0.000431303533715304 | AACUN_ASPA1 |
| NODE_13_length_81414_cov_33.042008_g4_i5 | 10.5848982044666 | 2.00112988566142e-11 | 4.35910641674014e-08 | AG45_MYCLE,6PGDH_GLUOX,ACRD_ECOLI,ACR_MYCTU |
| NODE_1302_length_7492_cov_102.596711_g16_i24 | 9.95740545772663 | 5.73554259885482e-07 | 0.000263476000418686 | ABGR_ECOLI |
| NODE_132_length_14297_cov_125.652208_g4_i48 | 11.1129142290899 | 1.02806343693862e-09 | 1.30986920319984e-06 | ACA11_ARATH |
| NODE_134_length_14240_cov_12.067975_g47_i0 | 10.4714634260852 | 1.96809355959952e-07 | 0.000103829235853622 | ACLDH_PSEAI,AEGA_ECOLI,3BHS_PIG |
| NODE_13413_length_3215_cov_748.032782_g1364_i3 | -11.9795269535717 | 3.37547308579046e-13 | 3.79898244228763e-09 | ABT3B_DANRE |
| NODE_136_length_14070_cov_13.372937_g49_i0 | 9.83362782499677 | 2.48981314785576e-07 | 0.000127372804733639 | ADPR_RHOER,5NTD_PSEAE,ABDH_CITK8,ACRS2_ALTAL |
| NODE_14_length_77286_cov_28.739176_g7_i0 | 11.6566463041542 | 1.77633130838562e-10 | 2.78958373471312e-07 | AFSQ1_STRCO,AAP1_YEAST,ACSA2_NOVHA,A2965_ARTBC,AFG2_YEAST,AFSQ2_STRCO,ADEC_LISMF,ADHI_CERS4,AFSQ2_STRCO,AGLJ_HALVD,ADRC2_ARAT |
| NODE_1401_length_7343_cov_16.089271_g16_i26 | 9.77866153083893 | 1.34801572717361e-06 | 0.000520164605854742 | ACLR_ACHOB,ADAB_ASPNG,ACOC_PSEPU |
| NODE_141_length_13928_cov_61.097077_g16_i10 | 9.19016397951263 | 1.22641253017483e-07 | 6.95942733929796e-05 | ACCO1_DICMU |
| NODE_14228_length_3118_cov_22.640722_g16_i46 | 7.99788962417331 | 4.20024465799637e-05 | 0.00903293379825411 | AER_ECOLI |
| NODE_14331_length_3106_cov_12.098582_g4893_i0 | 8.60145338425619 | 1.87706468651661e-05 | 0.00446318394898217 | AGL5_HALVD |
| NODE_1438_length_7278_cov_13.436919_g626_i0 | 9.56697566475207 | 1.62545292212163e-06 | 0.000593316675270431 | 128UP_DROME |
| NODE_1445_length_7269_cov_12.169956_g630_i0 | 9.46105361013296 | 1.94478109497234e-06 | 0.000694852792493609 | AAT_SULTO |
| NODE_146_length_13789_cov_10.634004_g52_i0 | 10.8490407011562 | 9.46332277039114e-08 | 5.65521469061038e-05 | ACT3_STRCO,ACSF2_MACFA,ADAT2_XENTR |
| NODE_147_length_13767_cov_12.164452_g53_i0 | 11.8691611078037 | 1.95625174654174e-09 | 2.44632903593465e-06 | AER_ECOLI,ACLY_HUMAN |
| NODE_1477_length_7212_cov_48.663118_g644_i1 | -7.33035398609527 | 6.41767938983116e-05 | 0.0128980075546583 | ACBD6_XENLA |
| NODE_1495_length_7179_cov_12.723755_g653_i0 | 9.90762316275277 | 8.73273108409571e-07 | 0.000365524807641599 | AFSQ2_STRCO |
| NODE_15165_length_3008_cov_55.530153_g139_i8 | -7.08885206810562 | 7.93026100379576e-05 | 0.0154326992813925 | AFAD_MOUSE |
| NODE_1564_length_7072_cov_12.178883_g677_i0 | 9.87367555665227 | 1.05687642963057e-06 | 0.00042735779365325 | AAEA_ECO8A,ACOC_BACSU,ACNR_CORGL |
| NODE_1597_length_7023_cov_12.964748_g694_i0 | 9.94605069347641 | 9.75693051486836e-07 | 0.00040174756329758 | ACLH_ASPOR,ADHI_CERS4 |
| NODE_16_length_74437_cov_42.698160_g4_i7 | 11.22174104958 | 6.47143648267575e-12 | 2.15762458510469e-08 | ACRD_ECOLI,4CL5_ORYSJ,ACSM3_PONAB,AGL2_BACTQ,ACD11_CAEEL,ACOX_SYNY3,AADH2_MAIZE,ACO13_MOUSE,ABNA_ASPCL,ACOCT_ECOLI,ACD11_CHIC |
| NODE_1620_length_6996_cov_13.324570_g705_i0 | 8.48915752025639 | 4.8650055448727e-06 | 0.00142836562797463 | AGLE_HALVD |
| NODE_16366_length_2881_cov_14.791311_g5490_i0 | 9.73410652319478 | 1.13771068267184e-06 | 0.000451925452820379 | AGLE_HALVD |
| NODE_164_length_13188_cov_85.686237_g4_i50 | 12.8745362635066 | 3.42016458847004e-10 | 4.91397604957883e-07 | ACLH_ASPOR |
| NODE_17_length_73955_cov_34.237703_g4_i8 | 6.99305304394285 | 1.76969485967066e-05 | 0.00423772888240569 | ACRD_ECOLI,ACRE_ECOLI,ABTIR_ACIB9,ADHR_BACSU,ACRD_ECOLI,ABAH3_ORYSJ,ACA11_ARATH,AGL5_HALVD,ACPY1_SACBS,ABR1_ASPFU,1CPX_DIRIM,1C |
| NODE_17075_length_2810_cov_1602.484107_g5710_i1 | -5.62884397062561 | 4.37405676519493e-06 | 0.00131862189839323 | ACRO_RAT |
| NODE_177_length_12837_cov_17.024522_g71_i0 | 5.88598268288654 | 0.000151098289497104 | 0.0267103803485876 | ACA10_ORYSJ,ADHR_BACSU,ADHR_BACSU |
| NODE_18_length_72403_cov_29.436361_g4_i9 | 8.63664841920648 | 3.84478059214457e-08 | 2.67660148274576e-05 | ACRD_ECOLI,ABTIR_ACIB9,AAEA_ECO7I,ADPR_LACLL,AFSQ1_STRCO |
| NODE_1810_length_6813_cov_11.519288_g791_i0 | 9.78335355064522 | 1.25801872344711e-06 | 0.000493904002075212 | AFSQ2_STRCO |
| NODE_183_length_12731_cov_18.072681_g71_i1 | 6.07233809807775 | 0.000112636629284633 | 0.0206128084074057 | ACA10_ORYSJ,ADHR_BACSU,ADHR_BACSU |
| NODE_1892_length_6738_cov_11.602251_g824_i0 | 10.8633215626416 | 5.63940568968061e-08 | 3.69726007196847e-05 | A7629_ARTBC |
| NODE_19_length_69203_cov_28.390308_g4_i10 | 15.3661632482723 | 1.19557090869073e-13 | 1.82628975154156e-09 | AGLUS_THESQ,ABFC_EMENI,6GAL_HYPRU,ABNB_ASPTN,ACRT_ASPA1,ADHA_KOMEU,ACA1_SCHPO,3BHD1_EGGLE |
| NODE_1902_length_6725_cov_13.326368_g828_i0 | 8.19513930358531 | 6.38785331117857e-06 | 0.00176786458359535 | ACL1_SCHPO |
| NODE_19229_length_2611_cov_10.134358_g6348_i0 | 8.22826102225872 | 3.31052996765999e-05 | 0.00742702550352637 | A9A1A_DANRE,ACAD8_DICDI |
| NODE_197_length_12464_cov_13.755952_g78_i0 | 11.8466292585686 | 5.64127406385033e-09 | 5.6021169850542e-06 | ACT22_STRCO,AAER_ECOLI,ACDC_ASPCN,ABGR_ECOLI |
| NODE_2_length_437557_cov_29.038669_g0_i1 | 10.741568035194 | 1.47438206990093e-08 | 1.15908032966428e-05 | AFSQ2_STRCO,ACOR_BACSU,AGL11_HALVD,ABAF_ACIBT,AGLA_RHIME,AAPA_BACSU,AGLB_THENE,ACOR_BACSU,AAAT_RABIT,AAAT_RABIT,ACCD_NOVAD,A |
| NODE_20_length_67517_cov_27.151904_g8_i0 | 11.7322870802354 | 1.67395893335954e-10 | 2.6914071155215e-07 | AB3F_ARATH,ACMB_GORST,ADCS_STRGR,ACMB_GORST,128UP_DROME,2NPD_CYBMR,AFG3_YEAST,8ODP_RAT |
| NODE_2003_length_6628_cov_12.364302_g867_i0 | 8.68932997985074 | 8.25518781825952e-06 | 0.00218416023187028 | AFSQ2_STRCO |
| NODE_20258_length_2524_cov_94262.546716_g6418_i1 | -21.5549047431653 | 3.51244299939252e-08 | 2.47071094648935e-05 | ADA1A_ORYLA |
| NODE_20663_length_2492_cov_1007.510128_g6539_i3 | 10.417046797779 | 2.42805634872923e-09 | 2.7790133748642e-06 | ACRO_RABIT |
| NODE_21_length_64405_cov_32.809939_g4_i11 | 7.16132914380148 | 2.34443443292048e-08 | 1.75905520429171e-05 | ADPR_RHOER,3BHS4_MOUSE,AADR_RHOPA,AFSQ1_STRCO,ACR_MYCTU,ACR_MYCTU |
| NODE_21192_length_2447_cov_15.165965_g6958_i0 | 8.46109087099462 | 2.59321135183024e-05 | 0.00591602622183758 | AAPAT_THET8 |
| NODE_2180_length_6462_cov_9.743465_g948_i0 | 9.42638519065367 | 2.78468344526082e-06 | 0.000908425621698418 | ADDA_LACLA |
| NODE_2193_length_6452_cov_21.565292_g16_i28 | 9.77304445507962 | 3.06932783117389e-07 | 0.000151288737068256 | ACA13_ARATH |
| NODE_22_length_63997_cov_32.363416_g4_i12 | 14.4673908352714 | 0.00012997224611813 | 0.0232189572377383 | ABKA_DICDI,ACEA_KOMXY,AGLM_HALVD,AGL12_HALVD,AGL12_HALVD,AGLE_HALVD,AGL14_HALVD,AGL12_HALVD,ADAA_BACSU,AGL11_HALVD,ADPR_LACLL |
| NODE_23_length_59380_cov_27.998061_g9_i0 | 13.0617309103182 | 3.37474557055932e-12 | 1.42431136805456e-08 | ACD10_HUMAN,AGL16_SULAC,ACEA_KOMXY,ADK_SCHCO,AGLI_HALVD,ACPH_PIG |
| NODE_2331_length_6338_cov_11.794254_g1011_i0 | 9.12556942382315 | 2.7618102559314e-07 | 0.000138157536393277 | AARE1_ORYSJ,AFSQ2_STRCO |
| NODE_2354_length_6324_cov_234.827228_g974_i1 | -8.49387527385859 | 9.16215655408382e-07 | 0.000379571845266364 | AEF1_DROME,AEF1_DROME,AEF1_DROME |
| NODE_2365_length_6317_cov_12.631967_g1028_i0 | 7.90416941406592 | 9.42911179631993e-06 | 0.00243957494782334 | ACCC1_POPTR,ADCS_SCHPO |
| NODE_237_length_11643_cov_11.977442_g96_i0 | 11.6253005643571 | 7.71524516712229e-09 | 6.9585028371797e-06 | ACSF2_RAT |
| NODE_2383_length_6298_cov_148.826827_g1030_i1 | -6.86364567427668 | 0.000199436619683399 | 0.0333355347870806 | 343L_IIV6 |
| NODE_24_length_58959_cov_29.525541_g4_i13 | 12.3846488520713 | 1.15671980811985e-11 | 2.78967768581133e-08 | AB1K7_ARATH,ABAQ_ACIBT,AFSQ1_STRCO,ABNA_ASPCL,ACI_VIBSJ,ABFB_MAGO7,AGL16_SULAC,AGL12_HALVD,AER_ECOLI |
| NODE_25_length_58089_cov_27.655561_g10_i0 | 10.3854563480352 | 5.51224693451551e-10 | 7.29864727439144e-07 | AAEA_YERE8,ADRX_YEAST,3O1D_COMTE,AFG1L_MOUSE,ACOC_BACSU,2KGR_GLUOX,ACLY_HUMAN,AAAH_CHLPN,3MG2_ECOLI,ABCH2_DICDI |
| NODE_2564_length_6169_cov_12.347441_g1105_i0 | 8.54219257609906 | 1.39789600786521e-05 | 0.00347048241246773 | AEPA_PECCC |
| NODE_26_length_52440_cov_37.777837_g4_i14 | 14.9557392899766 | 1.15620493231858e-12 | 6.50635055580077e-09 | ACSA_NOVAD,AER_ECOLI,AFSQ2_STRCO,AAEA_SERP5,AFSQ2_STRCO,AFSQ1_STRCO,AFSQ1_STRCO,ACCR_AGRFC,AFSQ1_STRCO,AFSQ1_STRCO |
| NODE_27_length_52152_cov_27.807446_g11_i0 | 12.5121507501963 | 9.06108956134211e-12 | 2.46958294693296e-08 | ACOB_BACSU,ACEA_KOMXY,AER_ECOLI,ABNB_EMENI,ACOA_BACSU,AGLK_RHIME,ACR1_ACIAD,AFLM_ASPPU,ADHR_BACSU |
| NODE_27137_length_2046_cov_11.409022_g8906_i0 | 6.91344854008874 | 0.000182777514701186 | 0.0309181393258695 | ACOB_BACSU,ACOA_BACSU |
| NODE_275_length_11152_cov_11.781749_g116_i0 | 11.5406918549041 | 1.22944557686869e-08 | 1.01162004866031e-05 | ADPP_METJA,ABAF_ACIBT,ACBC_ACTS5,ACCA_RHIWR |
| NODE_27500_length_2025_cov_10.482582_g9012_i0 | 7.84318555483599 | 8.15015407298872e-05 | 0.0157697307805382 | AAP2_NEUCR,AADH1_MALDO |
| NODE_28_length_51018_cov_33.238748_g4_i15 | 10.3969464285538 | 8.09759337240627e-09 | 7.01043955451091e-06 | AGL5_HALVD,AGLR_RHIME,AGLR_RHIME,AGLI_HALVD,A7629_ARTBC,AGLE_HALVD,ACLG_ASPOR,ACPH_PIG |
| NODE_28644_length_1963_cov_211.900000_g9387_i0 | 6.6933958977505 | 2.12325289044599e-05 | 0.00492711413010436 | 72A61_TRIFG |
| NODE_2960_length_5910_cov_12.734453_g1250_i0 | 9.79556818119171 | 1.40545110683532e-06 | 0.000539246036036225 | ADCS_STRVP |
| NODE_3_length_254142_cov_28.439696_g1_i0 | 10.5666005910456 | 2.89750040810942e-12 | 1.30441605039209e-08 | ACRA_ECOLI,AGLF_HALVD,6PGL_BACSU,ABC8B_MOUSE,AER_ECOLI,ACRS2_ALTAL,2KGR_VITVI,ADRG_PENRW,ADK_YEAST,AARE_ARATH,3HIDH_ARATH,AFS |
| NODE_30_length_50786_cov_27.425394_g13_i0 | 12.5036048209745 | 2.10761800496303e-11 | 4.44760089497324e-08 | AAU3_RHIME,ACEA_HALH5,AFR_RHIME,AFTS1_ALTAL,AER_ARATH,AAD_PHACH,AGLR_RHIME,AB8I_ARATH,3BHD2_EGGLE,ABCX_CYAPA,3BHDP_RUMGV |
| NODE_300_length_10964_cov_34.245065_g16_i14 | 9.98310963643525 | 7.99217906040748e-07 | 0.000339431363264903 | ACA7_ORYSJ,ADHR_BACSU |
| NODE_30306_length_1876_cov_12.348308_g9967_i0 | 8.00000805958451 | 5.64266325836928e-05 | 0.0116525310248061 | AACUQ_ASPA1 |
| NODE_30376_length_1873_cov_8.777778_g9989_i0 | 7.4984807307709 | 0.000174444665586474 | 0.0299742986710519 | AAT_SULTO |
| NODE_30508_length_1866_cov_10.811489_g10037_i0 | 7.94054949888841 | 6.08412220261346e-05 | 0.012487799516659 | AAD_PHACH,ACEA_PSEAE |
| NODE_30787_length_1852_cov_107.865655_g10126_i0 | 5.16306555658039 | 6.10270365850941e-05 | 0.0124879809894492 | AEF1_DROME |
| NODE_3086_length_5843_cov_12.165858_g1296_i0 | 9.58201894413769 | 2.06892680843886e-06 | 0.000727658799584682 | ACPH_PIG |
| NODE_32_length_47717_cov_40.808328_g4_i16 | 12.1028769576749 | 1.7584493885606e-11 | 3.95815234369068e-08 | A9A1B_DANRE,ADAS_DROME,ACOC_BACSU,ACOA_BACSU,ACOB_BACSU,ACT3_STRCO,3AHDP_RUMGV,ACTT3_ALTAL,ABR1_ASPFU |
| NODE_32465_length_1770_cov_914.790807_g10297_i1 | -6.57025665103383 | 0.000106918314003391 | 0.0197876710166077 | ACRB_ASPA1 |
| NODE_33_length_47184_cov_29.282609_g14_i0 | 12.7570186168112 | 5.56910625825056e-11 | 1.01640704704633e-07 | ACRD_ECOLI,AFG31_MOUSE,AAEA_SERP5,ACR3_ARATH,AB17I_ARATH,AACUM_ASPA1,ABHDA_BOVIN,AFRP_STRGR |
| NODE_33068_length_1744_cov_941.688211_g4047_i3 | -8.42646179591523 | 9.99610298856148e-07 | 0.000409101116734291 | A4_TAKRU |
| NODE_33167_length_1738_cov_98.823423_g10895_i1 | 6.86947394367042 | 0.000216464058784453 | 0.0360034112354595 | ADLO1_ARATH |
| NODE_33178_length_1737_cov_1657.323918_g10540_i4 | 7.78798832924633 | 1.42240625556339e-05 | 0.00351839742218625 | ACRO_HUMAN |
| NODE_3325_length_5719_cov_11.841658_g1378_i0 | 8.97022438999075 | 8.74359123610152e-07 | 0.000365524807641599 | ACTT3_ALTAL |
| NODE_333_length_10712_cov_12.482846_g143_i0 | 10.4408754770701 | 7.50010408219659e-08 | 4.67699309898071e-05 | ADHR_BACSU,ACDB_BACSU,ACOB_BACSU,ACOA_BACSU |
| NODE_34_length_46982_cov_36.291330_g4_i17 | 12.5741095605625 | 7.0293420317947e-12 | 2.15762458510469e-08 | 2KGR_VITVI,ABC1_AJECA,AAT_MUSP7,AES1_SCHPO,AFLM_ASPPU,4HYPE_BRUSU |
| NODE_3419_length_5672_cov_11.138596_g1417_i0 | 7.22682358115258 | 4.11906320765578e-05 | 0.00894379743686751 | 1A1D_THEMA,ACCD_NOVAD |
| NODE_3439_length_5658_cov_11.669114_g1420_i0 | 9.5947773049345 | 1.56896118051453e-06 | 0.000578955249168226 | AAAD_RABIT |
| NODE_34792_length_1667_cov_387.534504_g9446_i3 | 9.59792390683979 | 9.07275407067146e-09 | 7.65831171105378e-06 | ACPH1_APIME |
| NODE_35_length_46449_cov_31.008625_g4_i18 | 13.9716500200041 | 6.61058492681474e-12 | 2.15762458510469e-08 | ABFB_MAGO7,AFSQ2_STRCO,AADH2_PEA,ABAQ_ACIBT,AGLR_RHIME,AGLR_RHIME,ACTT6_ALTAL,AGL2_BACTQ,A622L_TOBAC,ADAT2_XENTR,AFSQ1_STRCO, |
| NODE_356_length_10488_cov_39.188190_g4_i58 | 12.8588614854635 | 2.42847214223101e-10 | 3.69587825129235e-07 | ACNR_CORGL |
| NODE_36_length_45573_cov_39.489780_g4_i19 | 12.7027012003509 | 3.37372616629664e-10 | 4.91397604957883e-07 | ACDAS_XANCP,128UP_DROME,3BHS_MYCTU,AFSQ2_STRCO,AFSQ2_STRCO,AER_ECOLI,AFSQ1_STRCO,ACLR_ACHOB |
| NODE_36511_length_1590_cov_9.760053_g12243_i0 | 8.55776395150165 | 1.5166988973517e-05 | 0.00371085663552049 | A7629_ARTBC |
| NODE_37_length_45362_cov_32.858487_g4_i20 | 11.0991915763149 | 7.53948590856181e-11 | 1.33980632745622e-07 | 11SB_CUCMA,6PGL_SODGM,ACPH_RAT,AB25B_ARATH,ADAB_BACSU |
| NODE_37708_length_1535_cov_44.381669_g12721_i0 | -6.8180128703879 | 0.000175974513627087 | 0.0300586082242757 | A4GAT_RAT |
| NODE_37828_length_1530_cov_9.492793_g12768_i0 | 7.47520512390846 | 0.000173084339091861 | 0.0298164266586612 | AARE1_ORYSJ |
| NODE_38_length_44095_cov_30.947140_g4_i21 | 28.0985102024048 | 6.37366321579035e-13 | 4.70373912288712e-09 | ACT22_STRCO,ACCO1_DICMU,ACDA1_PYRFU,ACSL5_RAT |
| NODE_3828_length_5453_cov_10.933457_g1559_i0 | 9.22288737698085 | 4.33380957899021e-06 | 0.00131825897860383 | AER_ECOLI |
| NODE_388_length_10232_cov_11.711684_g165_i0 | 9.96126084882749 | 6.73565251910588e-07 | 0.000296052885894764 | 6PGDH_GLUOX,AFSQ1_STRCO |
| NODE_39_length_43124_cov_83.650298_g4_i22 | 8.07875849783757 | 2.25342381555449e-09 | 2.66963514766252e-06 | ADHB_BACSU,ADHB_BACSU,ABC1_MYCTU,ABGR_ECOLI,ADRC1_ARATH,ADHL_GADMO,ADHR_BACSU |
| NODE_4_length_227115_cov_27.563847_g2_i0 | 11.3832841484363 | 6.96561296482515e-13 | 4.70373912288712e-09 | AGLUS_ARCFU,AAAT_BOVIN,ABVS_ASPBC,AGLR_RHIME,ACAC_DICDI,AGGR_ECOLX,AGLD_HALVD,3BHS_PIG,ADPR_RHOER,AFLW_ASPPU,ABCA4_MOUSE,AC |
| NODE_40_length_43124_cov_83.220274_g4_i23 | 11.033768372891 | 4.8552572341296e-11 | 9.10738362517511e-08 | ADHB_BACSU,ADHB_BACSU,ABC1_MYCTU,ABGR_ECOLI,ADRC1_ARATH,ADHL_GADMO,ADHR_BACSU |
| NODE_40306_length_1430_cov_17908.750184_g313_i181 | -12.95891317522 | 2.68597158187313e-15 | 1.12897076471271e-10 | ABRU_DROME |
| NODE_409_length_10092_cov_11.505440_g179_i0 | 9.7877633444688 | 1.11971714204048e-06 | 0.000447409817560412 | ACOA_CUPNH,ACADL_PIG |
| NODE_41_length_42709_cov_33.859508_g4_i24 | 12.9135694022463 | 5.83277863080807e-12 | 2.15762458510469e-08 | ACOC_BACSU,ACYLL_CAUVN,AGLF_HALVD,ACO11_MOUSE |
| NODE_42_length_42197_cov_30.632632_g4_i25 | 11.0024260413289 | 5.81115033534743e-09 | 5.60593371207631e-06 | AGLA_RHIME,AGLA_RHIME,AFSQ2_STRCO,AGAA2_SHESA,AGAS_ECOLI,AGL9_HALVD,AGLR_RHIME,ACOT8_HUMAN,AADR_RHOPA,AFSQ1_STRCO,AGAL_BACT |
| NODE_43_length_42007_cov_38.359851_g4_i26 | 11.8081727779389 | 3.70222498480919e-11 | 7.35305437571161e-08 | ABD12_MACFA,5NTD_PSEAE,A57_VACCC |
| NODE_44_length_32067_cov_34.639714_g4_i27 | 14.3591196564704 | 5.93256286715029e-12 | 2.15762458510469e-08 | AFSQ2_STRCO,AG45_MYCLE,25DK1_PSEPU,ABCA9_DICDI,AAPP_RHIL3,ABCF3_ENCCU |
| NODE_44176_length_1286_cov_15.437758_g15497_i0 | 7.63654843697875 | 9.91746332861552e-05 | 0.018602957323743 | AB8I_ARATH |
| NODE_46_length_31921_cov_42.270535_g4_i29 | 12.850628052483 | 1.02222919426277e-10 | 1.7699767443635e-07 | AGL5_HALVD |
| NODE_4659_length_5111_cov_11.934895_g1857_i0 | 10.4705542242755 | 1.61063592052865e-07 | 8.96167330003103e-05 | AARP_PROST |
| NODE_47_length_31537_cov_93.347254_g4_i30 | 9.59173189813674 | 1.1518039206501e-10 | 1.9444753788415e-07 | ACRD_ECOLI,ADHR_BACSU |
| NODE_4902_length_5018_cov_13.194742_g1949_i0 | 7.3036914998343 | 5.59764137511413e-05 | 0.011595016158856 | ACRD_ECOLI |
| NODE_5_length_199014_cov_28.510991_g3_i0 | 12.6339713132854 | 1.35224629156909e-13 | 1.82628975154156e-09 | ACXA_XANP2,ADHB_BACSU,ACR3_ALKMQ,ADRC1_ARATH,ACOB_BACSU,2KGR_VITVI,ACOC_BACSU,3AHD_EGGLE,AFSQ1_STRCO,ABCH2_DICDI,AFSQ1_STRC |
| NODE_5185_length_4905_cov_12.887003_g2049_i0 | 8.30217394245286 | 1.8843506684773e-05 | 0.00446478708564685 | ADAB_ASPNG |
| NODE_5191_length_4901_cov_14.123239_g2053_i0 | 9.64927971922259 | 1.41959944615081e-06 | 0.000541597239546169 | AFSQ2_STRCO |
| NODE_53_length_27289_cov_33.765175_g4_i33 | 14.0483367495165 | 3.72036392707478e-05 | 0.00815677711907487 | ADHC2_MYCS2,235L_IIV6,AB1I_ARATH |
| NODE_53458_length_1006_cov_608.711683_g1364_i13 | -6.49487093389562 | 5.20531092674298e-05 | 0.0108824840947709 | ABAP1_ARATH |
| NODE_54_length_26230_cov_31.687770_g4_i34 | 11.8009045249889 | 3.11248744339765e-09 | 3.23353926273472e-06 | ADAB_ASPNG,ACON3_ASPFU,ACON3_ASPFU,AER_ECOLI |
| NODE_54006_length_992_cov_2225.140370_g14771_i1 | -8.66306401992816 | 3.72796437607581e-07 | 0.000178540410204005 | ACY1_PIG |
| NODE_5402_length_4842_cov_12.112812_g2122_i0 | 8.56782636551434 | 1.97001259561225e-06 | 0.000700163213455285 | AATC_RHIME |
| NODE_542_length_9325_cov_13.306312_g242_i0 | 9.31296177292679 | 6.78222277383583e-07 | 0.000296052885894764 | ACRD_ECOLI,AAPA_BACSU,AFSQ2_STRCO,AFSQ1_STRCO |
| NODE_5437_length_4831_cov_12.602984_g2131_i0 | 10.4483251127051 | 1.14713979216065e-07 | 6.56475049873087e-05 | ACCA1_MYCTU,AAEA_PECAS |
| NODE_5489_length_4812_cov_11.884786_g2146_i0 | 9.24841870032814 | 3.96552602684957e-06 | 0.00122836716303256 | AFR_ENSAD,AFTS1_ALTAL |
| NODE_55_length_25046_cov_29.210027_g18_i0 | 10.9507331598233 | 2.61841162805157e-09 | 2.94693500698445e-06 | AFSQ1_STRCO,ACBP4_ORYSJ,AFSQ2_STRCO,ACOR_BACSU |
| NODE_556_length_9267_cov_12.315532_g248_i0 | 9.88117532655839 | 7.40927419499242e-07 | 0.000319101641460802 | 1CPX_DIRIM,ABGR_ECOLI |
| NODE_5587_length_4778_cov_13.244633_g2189_i0 | 8.35822837445249 | 7.25463027278361e-06 | 0.00195175566956387 | AFLM_ASPPU |
| NODE_56_length_24230_cov_39.107795_g4_i35 | 11.6952135695602 | 2.22540083039468e-09 | 2.66963514766252e-06 | ABAB_STRAT,AADR_RHOPA,AADR_RHOPA,AFG2_SCHPO |
| NODE_57_length_24172_cov_99.654923_g4_i36 | 11.6111861338614 | 2.95258465575326e-09 | 3.16479581958263e-06 | ACRF_ECOLI,AFSQ2_STRCO |
| NODE_58_length_23630_cov_32.048266_g4_i37 | 10.5163529701139 | 5.19193083676232e-08 | 3.4712941143058e-05 | ACR1_ACIAD,AFLM_ASPPU,ABGR_ECOLI |
| NODE_59_length_23481_cov_28.263713_g19_i0 | 13.8434506012659 | 3.12234272344715e-11 | 6.38925937663451e-08 | AGL16_SULAC,2NPD_NEUCR,ACPH_RAT,ACPS_SPHAL |
| NODE_591_length_9129_cov_11.832818_g260_i0 | 10.3863989171144 | 5.41921035762485e-08 | 3.58772977480089e-05 | A9A1B_DANRE,ABHDB_BOVIN,ACT3_STRCO |
| NODE_5953_length_4669_cov_15.709530_g2291_i0 | 5.04049615575347 | 0.000277801710333662 | 0.0446652235605036 | 6PGDH_GLUOX |
| NODE_6_length_170876_cov_29.570745_g4_i0 | 11.809369559849 | 1.4612590928465e-11 | 3.40261738005995e-08 | AGL11_HALVD,AGL2_BACTQ,AADR_RHOPA,AADR_RHOPA,AFSQ1_STRCO,AB7B_ARATH,3BHD1_EGGLE,ACA6_ORYSJ,ACVA_ACRCH,AER_ECOLI,438L_IIV6,AARD |
| NODE_61_length_21781_cov_28.135664_g20_i0 | 11.6413388785837 | 2.84699310768796e-09 | 3.10083468670891e-06 | AGLD_HALVD,ABAB_STRAT,ACLR_ACHOB |
| NODE_623_length_8967_cov_17.354509_g272_i0 | 11.8005822775635 | 2.00837983557038e-09 | 2.4658522461163e-06 | AEEP_BACSU |
| NODE_625_length_8961_cov_11.840459_g273_i0 | 9.07247974589551 | 1.45569217332476e-06 | 0.000546111006001526 | ADPR_LACLL |
| NODE_6269_length_4586_cov_157.975183_g1364_i2 | -6.34345450098351 | 7.8596060359542e-06 | 0.00208954124565321 | ABAP1_ARATH |
| NODE_6299_length_4579_cov_10.438748_g2407_i0 | 9.08047021245543 | 5.65823033705312e-06 | 0.00161551727353708 | ACA9_ORYSJ |
| NODE_6332_length_4570_cov_11.695352_g2421_i0 | 10.2121963482424 | 2.13503329871098e-07 | 0.000110903483534888 | 2KGR_VITVI,AAAD_RABIT |
| NODE_6395_length_4555_cov_11.429273_g2441_i0 | 9.21628798270122 | 3.98601593521883e-06 | 0.00122907618298382 | 3O1D_MYCS2 |
| NODE_64_length_21073_cov_51.547000_g4_i38 | 10.8479209125677 | 2.92518119693065e-08 | 2.12399608458422e-05 | AFSQ1_STRCO |
| NODE_6526_length_4513_cov_13.664640_g2482_i0 | 9.14743095122113 | 5.12436208855231e-06 | 0.00148858271998232 | AFSQ2_STRCO,ABCF3_ENCCU,AFSQ1_STRCO |
| NODE_699_length_8699_cov_11.864364_g308_i0 | 10.051426810337 | 7.70890847350624e-07 | 0.000329472893290462 | ACSA_NOVAD |
| NODE_7_length_110398_cov_31.382388_g4_i1 | 12.8422874762561 | 1.70160016313723e-12 | 8.83889660125624e-09 | ACNR_MYCTU,AEPA_PECCC,ACRB_ECOLI,ACRT_ASPA1,AAEA_ECOHS,AGL9_HALVD,72A61_PARPY,AGL16_SULAC,ACDC_ASPCN,ACRA_ECOLI,AGL5_HALVD,AD |
| NODE_7226_length_4333_cov_13.970423_g2704_i0 | 10.375687237912 | 2.76200500726993e-07 | 0.000138157536393277 | ABCA5_DICDI |
| NODE_74_length_19613_cov_27.783572_g24_i0 | 13.5902406381491 | 4.10119194451992e-11 | 7.91272256084404e-08 | ADH1_GEOSE,ADHS_AQUAE,ACCA3_MYCTU,3DHQ_TALSN,3HAPM_CUPPJ,ACCC_HAEIN |
| NODE_7473_length_4268_cov_11.312992_g2774_i0 | 9.01450985996498 | 6.14474957767131e-06 | 0.00171732245343143 | AEPA_PECCC,AFSQ1_STRCO |
| NODE_78_length_18804_cov_19.526987_g26_i0 | 6.96988979767674 | 2.03164202072065e-06 | 0.000718286504582326 | ABGR_ECOLI,AAER_ECOLI,A622L_TOBAC,3AHDP_RUMGV,AAMHY_RHILO,ADH_SULTO,ABGR_ECOLI |
| NODE_8_length_106877_cov_30.445976_g4_i2 | 12.3666636106385 | 1.99578508041671e-12 | 9.62652677931281e-09 | ACRS2_ALTAL,ABCH2_DICDI,AGD2_ARATH,ADCS_STRGR,AFVA_ASPFN,ACLR_ACHOB,ACTT3_ALTAL,3HBH1_PSEAC,AATC_RHIME,AAPAT_CERS1,AAT_BACY2,A |
| NODE_810_length_8417_cov_25.688279_g363_i0 | 10.0128131845786 | 1.03797154614754e-07 | 6.04242608346992e-05 | AFSQ1_STRCO,AFSQ2_STRCO |
| NODE_8214_length_4095_cov_12.013426_g3031_i0 | 9.11470671921092 | 4.77767073782014e-06 | 0.0014150287262435 | AGL9_HALVD |
| NODE_83_length_18023_cov_80.397827_g4_i42 | 13.2177780001198 | 0.000221447321978304 | 0.0365620898742076 | 3AHDP_RUMGV |
| NODE_8335_length_4068_cov_14.209512_g3070_i0 | 8.47599261395652 | 9.38729329963129e-06 | 0.00243809669975962 | 3HAPM_CUPPJ |
| NODE_84_length_17956_cov_22.383884_g17_i2 | 8.28800147071204 | 7.75175770600506e-09 | 6.9585028371797e-06 | AFSQ1_STRCO |
| NODE_863_length_8267_cov_11.765926_g388_i0 | 11.1194911997697 | 2.00526679361503e-08 | 1.53876881862768e-05 | AFSQ2_STRCO |
| NODE_87_length_17732_cov_13.364630_g29_i0 | 9.67679746738986 | 1.72560323920351e-07 | 9.39730125297858e-05 | ACD17_MYCTU,ACDSB_BOVIN,ADHS_METM6,ACTT3_ALTAL,5FCL_HAEIN,ACSF2_MOUSE,ACMB_GORST |
| NODE_8734_length_3985_cov_13.176125_g3194_i0 | 10.1511686541308 | 4.98915505124374e-07 | 0.000233496646647901 | ACT3_STRCO,ACDA1_PYRFU |
| NODE_888_length_8211_cov_12.218481_g397_i0 | 6.93537007524658 | 6.34992564880899e-06 | 0.00176459991445586 | AGL5_HALVD |
| NODE_9_length_101437_cov_27.937917_g4_i3 | 11.9655439325988 | 1.06573514962063e-11 | 2.66544308087341e-08 | ACMSD_HUMAN,AER_ECOLI,ACES_BUNFA,ADEC_LISMO,ACT22_STRCO,AFR_ENSAD,AFSQ1_STRCO,ADI1_USTMD,ABAB_STRAT,AAER_ECOLI,3AHDP_RUMGV,A |
| NODE_9447_length_3843_cov_8.872414_g3427_i0 | 8.8935756476559 | 7.3499975466136e-06 | 0.00196178116335069 | ACY1_MOUSE,AAAT_RABIT,ACNR_CORDI,ABSAA_ALCSP |
| NODE_95_length_16604_cov_36.197689_g4_i44 | 26.6421384874501 | 9.14281093373476e-12 | 2.46958294693296e-08 | ADC2_ORYSJ,ACSF2_HUMAN |
| NODE_9794_length_3777_cov_28.846652_g16_i40 | 6.65109148472049 | 3.61968868476306e-05 | 0.00796190024445212 | ADHR_BACSU |

Supplementary data 4C – Differentially expressed transcripts in adults of *Aedes koreicus*

| ID | log2FoldChange | pvalue | padj | Gene symbol |
|---|---|---|---|---|
| NODE_10480_length_2906_cov_71.230498_g3643_i1 | -21.8605376304313 | 3.68033325178351e-10 | 3.28658569951321e-06 | 7UP2_DROME |
| NODE_533_length_8398_cov_514.075435_g116_i4 | -21.8820558318018 | 3.52083712872002e-10 | 3.28658569951321e-06 | ACAC_CHICK |
| NODE_6590_length_3735_cov_275.428181_g2874_i1 | 10.4584483777863 | 7.72636304300686e-05 | 0.0313474501874027 | ABL1_CAEEL |
| NODE_16708_length_2109_cov_1627.423379_g5473_i5 | 10.8013272668994 | 2.90804354237122e-05 | 0.0121934630758011 | 5NTD_LUTLO |
| NODE_1077_length_6981_cov_644.463086_g498_i4 | 11.2220672644271 | 1.17782576973221e-05 | 0.00502270016434781 | ABLM3_HUMAN |
| NODE_31194_length_1121_cov_635.579198_g11149_i1 | 12.0206233586853 | 9.65640222394795e-06 | 0.00413545741909614 | ACH2_LONAC |
| NODE_8607_length_3249_cov_72.238665_g3160_i2 | 19.0854559976967 | 6.3566732264039e-08 | 2.7817528856082e-05 | AGAP9_HUMAN |
| NODE_31751_length_1095_cov_259.536204_g10294_i3 | 19.0854560379619 | 6.35667282315163e-08 | 2.7817528856082e-05 | ACR1_ACIAD |
| NODE_40939_length_780_cov_12.683168_g21471_i0 | 19.0854560379619 | 6.35667282315163e-08 | 2.7817528856082e-05 | AATNT_GEOSL |
| NODE_38877_length_836_cov_222.389253_g9367_i3 | 19.0854560468253 | 6.35667273493707e-08 | 2.7817528856082e-05 | ADNT1_ARATH |
| NODE_60299_length_506_cov_17.364896_g39784_i0 | 19.105560113943 | 6.17761113973928e-08 | 2.7817528856082e-05 | ACC1_CAEEL |
| NODE_84301_length_380_cov_201.475570_g63690_i0 | 20.5018391814119 | 6.23124028726323e-09 | 3.28658569951321e-06 | ACE_DROME |
| NODE_54601_length_559_cov_22.798354_g34206_i0 | 20.5222489110209 | 5.99977249827294e-09 | 3.28658569951321e-06 | AAMP_PONAB |
| NODE_35408_length_946_cov_12.240550_g17054_i0 | 20.5435730460876 | 5.76892576871195e-09 | 3.28658569951321e-06 | ACRO_HUMAN |
| NODE_9750_length_3040_cov_67.627570_g3997_i2 | 20.5435730460876 | 5.76892576871195e-09 | 3.28658569951321e-06 | 3BP5L_XENLA |
| NODE_39640_length_814_cov_16.171390_g20382_i0 | 20.5435731515257 | 5.76892473658706e-09 | 3.28658569951321e-06 | ACRO_SHEEP |
| NODE_30584_length_1151_cov_7.264378_g13822_i0 | 20.5435732263679 | 5.76892400394648e-09 | 3.28658569951321e-06 | ACRO_RAT |
| NODE_49578_length_622_cov_12.313297_g29378_i0 | 20.5618345064206 | 5.57677108644488e-09 | 3.28658569951321e-06 | ACRO_MOUSE |
| NODE_35685_length_936_cov_9.242178_g17267_i0 | 20.5618345193388 | 5.57677096407857e-09 | 3.28658569951321e-06 | A2GL_HUMAN |
| NODE_5059_length_4190_cov_22.617197_g2288_i0 | 20.5618345193388 | 5.57677096407857e-09 | 3.28658569951321e-06 | A2ML1_HUMAN |
| NODE_47385_length_653_cov_18.122414_g27295_i0 | 20.561834590913 | 5.57677028609683e-09 | 3.28658569951321e-06 | ACL6A_MOUSE |
| NODE_28763_length_1241_cov_9.779966_g12739_i0 | 20.5802032771692 | 5.39089553665435e-09 | 3.28658569951321e-06 | AEF1_DROME |
| NODE_16060_length_2175_cov_155.135585_g5723_i1 | 20.5802033427767 | 5.39089493530562e-09 | 3.28658569951321e-06 | AEF1_DROME |
| NODE_36511_length_908_cov_8.196407_g17895_i0 | 20.6163371183092 | 5.04496016624459e-09 | 3.28658569951321e-06 | A1131_ARTBC |
| NODE_37385_length_880_cov_10.355638_g18568_i0 | 20.6347705075207 | 4.87838373346377e-09 | 3.28658569951321e-06 | AEF1_DROME |
| NODE_60537_length_504_cov_29.814385_g40019_i0 | 20.634770537066 | 4.87838348769115e-09 | 3.28658569951321e-06 | ACACB_HUMAN |
| NODE_38447_length_849_cov_10.320876_g19409_i0 | 20.6347705800854 | 4.87838312983298e-09 | 3.28658569951321e-06 | 343L_IIV6 |
| NODE_31161_length_1123_cov_319.752381_g4940_i4 | 20.6522898792913 | 4.72528931898561e-09 | 3.28658569951321e-06 | A1AT_BOVIN |
| NODE_16830_length_2098_cov_162.321481_g3371_i1 | 20.6522901873437 | 4.72528680900768e-09 | 3.28658569951321e-06 | AGLB1_PYRFU |
| NODE_6975_length_3633_cov_353.192135_g586_i2 | 20.6522902630698 | 4.72528619828794e-09 | 3.28658569951321e-06 | 343L_IIV6 |
| NODE_19229_length_1881_cov_85.408186_g6183_i1 | 20.6671712524319 | 4.59823770977125e-09 | 3.28658569951321e-06 | A2GL_HUMAN |
| NODE_22512_length_1637_cov_190.627877_g9493_i1 | 20.6671712524319 | 4.59823770977125e-09 | 3.28658569951321e-06 | A622L_TOBAC |
| NODE_53992_length_566_cov_65.338742_g33626_i0 | 20.6671713914892 | 4.59823661759119e-09 | 3.28658569951321e-06 | AEF1_DROME |
| NODE_25030_length_1467_cov_10.277618_g10732_i0 | 20.686090520015 | 4.44439049077688e-09 | 3.28658569951321e-06 | AGL16_SULAC |

Supplementary data 4D – Differentially expressed transcripts in larvae of *Aedes koreicus*

| ID | log2FoldChange | pvalue | padj | Gene symbol |
|---|---|---|---|---|
| NODE_12687_length_2577_cov_52.490415_g5378_i0 | 6.690872860001 | 4.74391128841173e-05 | 0.0302612465256305 | AAP1_YEAST |
| NODE_15490_length_2231_cov_527.944856_g6513_i1 | 8.58078903702675 | 1.89370515103071e-05 | 0.0142020209144258 | ACOD1_MOUSE |
| NODE_21779_length_1686_cov_117.908246_g9158_i0 | 5.19542376090884 | 2.49477077064259e-05 | 0.0179808173322535 | A1345_ARTBC |
| NODE_29761_length_1189_cov_209.447133_g2550_i1 | 20.7533181174215 | 1.11222778598797e-07 | 0.000158270042897923 | ABCC8_DICDI |
| NODE_3552_length_4829_cov_241.814550_g527_i2 | -9.35835022185468 | 2.53852231379873e-07 | 0.000343610665485093 | AEF1_DROME |
| NODE_38031_length_860_cov_137.978399_g2329_i5 | 8.8540556995651 | 1.31909757174171e-06 | 0.00149399914161122 | ABCC8_DICDI |
| NODE_4224_length_4510_cov_221.697769_g1388_i2 | 20.7533181327582 | 1.11222776206333e-07 | 0.000158270042897923 | ACUC_STAAW |
| NODE_4656_length_4337_cov_415.782833_g749_i5 | -8.82835537460536 | 7.69941285410082e-07 | 0.000909136840774539 | AAPK2_CAEEL |
| NODE_6292_length_3805_cov_394.972401_g2514_i1 | 8.45251528821141 | 2.20145978264381e-05 | 0.0160755807312347 | 4HBCL_RHOPA |
| NODE_861_length_7384_cov_244.004240_g212_i2 | 20.7533182013147 | 1.11222765511911e-07 | 0.000158270042897923 | ABL2_MOUSE |
| NODE_874_length_7352_cov_170.563676_g406_i0 | -8.11163782714566 | 9.00481799139302e-06 | 0.00780844350106778 | AFOG_EMENI, ACTS3_ALTAL |
| NODE_9232_length_3127_cov_445.594957_g2239_i2 | 20.774553208796 | 1.07917531498577e-07 | 0.000158270042897923 | AEF1_DROME |
