## Supplementary table 5 for "De-novo genome assembly of the invasive mosquito species *Aedes japonicus* and *Aedes koreicus*"

Table S1. GO enrichment analysis of *Aedes koreicus* larvae.

| Name | p-value | Adjusted p-value | Odds Ratio | Combined score | Category |
|---|---|---|---|---|---|
| ATP-activated inward rectifier potassium channel activity (GO:0015272) | 0.002997 | 0.04453 | 444.11 | 2580.41 | Molecular Function |
| inward rectifier potassium channel activity (GO:0005242) | 0.01243 | 0.04453 | 92.44 | 405.55 | Molecular Function |
| actin monomer binding (GO:0003785) | 0.01243 | 0.04453 | 92.44 | 405.55 | Molecular Function |
| potassium ion transmembrane transporter activity (GO:0015079) | 0.01589 | 0.04453 | 71.54 | 296.32 | Molecular Function |
| non-membrane spanning protein tyrosine kinase activity (GO:0004715) | 0.01737 | 0.04453 | 65.22 | 264.33 | Molecular Function |
| phosphotyrosine residue binding (GO:0001784) | 0.01835 | 0.04453 | 61.59 | 246.23 | Molecular Function |
| protein phosphorylated amino acid binding (GO:0045309) | 0.02081 | 0.04453 | 54.06 | 209.35 | Molecular Function |
| manganese ion binding (GO:0030145) | 0.02375 | 0.04453 | 47.15 | 176.34 | Molecular Function |
| cation-transporting ATPase complex (GO:0090533) | 0.007973 | 0.03986 | 147.96 | 714.91 | Cellular Component |

Table S2. GO enrichment analysis of *Aedes koreicus* adults.

| Name | p-value | Adjusted p-value | Odds Ratio | Combined score | Category |
|---|---|---|---|---|---|
| fucose binding (GO:0042806) | 0.005488 | 0.04060 | 237.79 | 1237.70 | Molecular Function |
| CoA carboxylase activity GO:0016421) | 0.006583 | 0.04060 | 190.22 | 955.53 | Molecular Function |

Table S3. GO enrichment analysis of *Aedes japonicus* adults.

| Name | p-value | Adjusted p-value | Odds Ratio | Combined score | Category |
|---|---|---|---|---|---|
| purine nucleoside bisphosphate metabolic process (GO:0034032) | 1.362e-7 | 0.00004289 | 22.20 | 350.91 | Biological Process |
| ribonucleoside bisphosphate metabolic process (GO:0033875) | 1.362e-7 | 0.00004289 | 22.20 | 350.91 | Biological Process |
| acyl-CoA metabolic process (GO:0006637) | 8.603e-6 | 0.001807 | 14.67 | 171.07 | Biological Process |
| acyl-CoA dehydrogenase activity (GO:0003995) | 1.842e-4 | 0.01934 | 37.51 | 322.54 | Molecular Function |
