## Supplementary table 6 for "De-novo genome assembly of the invasive mosquito species *Aedes japonicus* and *Aedes koreicus*"

Table 1. Reads count of whole genome sequencing

| Nova seq 6000 | | |
|---|---|---|
| Library ID | Species | Reads |
| 80133_ID2319_1-Aejap | *Ae japonicus* | 571.666.400 |
| 80134_ID2319_2-Aekor | *Ae koreicus* | 717.967.340 |

| Oxford Nanopore | | |
|---|---|---|
| Library ID | Species | Reads |
| ID2319_1-Aejap | *Ae japonicus* | 2.476.846 |
| ID2319_2-Aekor | *Ae koreicus* | 4.369.744 |

Table 2 . Reads count of RNA sequencing (Nova Seq 6000)

| Library_ID | Reads (M) | Species | Life stage | Temperature reared (°C) |
|---|---|---|---|---|
| ID2684_1-Sample1-P1-A01 | 39.12 | *Ae koreicus* | Larvae | 15 |
| ID2684_2-Sample2-P1-B01 | 57.39 | *Ae koreicus* | Larvae | 15 |
| ID2684_3-Sample3-P1-C01 | 55.57 | *Ae koreicus* | Larvae | 15 |
| ID2684_4-Sample4-P1-D01 | 51.02 | *Ae japonicus* | Larvae | 15 |
| ID2684_5-Sample5-P1-E01 | 74.82 | *Ae japonicus* | Larvae | 15 |
| ID2684_6-Sample6-P1-F01 | 60.75 | *Ae japonicus* | Larvae | 15 |
| ID2684_7-Sample7-P1-G01 | 50.14 | *Ae koreicus* | Larvae | 28 |
| ID2684_8-Sample8-P1-H01 | 41.72 | *Ae koreicus* | Larvae | 28 |
| ID2684_9-Sample9-P1-A02 | 42.40 | *Ae koreicus* | Larvae | 28 |
| ID2684_10-Sample10-P1-B02 | 48.62 | *Ae japonicus* | Larvae | 28 |
| ID2684_11-Sample11-P1-C02 | 61.27 | *Ae japonicus* | Larvae | 28 |
| ID2684_12-Sample12-P1-D02 | 159.07 | *Ae japonicus* | Larvae | 28 |
| ID2684_13-Sample13-P1-E02 | 75.05 | *Ae koreicus* | Adults | 15 |
| ID2684_14-Sample14-P1-F02 | 40.04 | *Ae koreicus* | Adults | 15 |
| ID2684_15-Sample15-P1-G02 | 44.66 | *Ae koreicus* | Adults | 15 |
| ID2684_16-Sample16-P1-H02 | 40.34 | *Ae koreicus* | Adults | 28 |
| ID2684_17-Sample17-P1-A03 | 44.97 | *Ae koreicus* | Adults | 28 |
| ID2684_18-Sample18-P1-B03 | 49.55 | *Ae koreicus* | Adults | 28 |
| ID2684_19-Sample19-P1-C03 | 45.23 | *Ae japonicus* | Adults | 15 |
| ID2684_20-Sample20-P1-D03 | 48.63 | *Ae japonicus* | Adults | 15 |
| ID2684_21-Sample21-P1-E03 | 165.37 | *Ae japonicus* | Adults | 15 |
| ID2684_22-Sample22-P1-F03 | 39.67 | *Ae japonicus* | Adults | 28 |
| ID2684_23-Sample23-P1-G03 | 61.71 | *Ae japonicus* | Adults | 28 |
| ID2684_24-Sample24-P1-H03 | 41.65 | *Ae japonicus* | Adults | 28 |
